# Estrous cycle stage gates the effect of stress on reward learning

**DOI:** 10.1101/2025.03.23.644830

**Authors:** Morgan P. Johnston, Brandon I. Garcia-Castañeda, Leonor G. Cedillo, Sachi K. Patel, Victoria S. Vargas, Matthew J. Wanat

## Abstract

Stress produces transient physiological responses that lead to long-lasting changes in cue-driven behavior. In particular, a single exposure to stress facilitates reward learning in male rats. Since stress can produce distinct behavioral phenotypes between males and females, it is critical to additionally determine how stress impacts reward learning in females. To address this, female rats were exposed to restraint stress immediately prior to training on an appetitive Pavlovian conditioning task with food rewards. Females were categorized based on their estrous cycle stage on the first day of Pavlovian conditioning. A single exposure to stress enhanced conditioned responding in non-estrus females but suppressed conditioned responding in estrus females. Therefore, a single stress experience produced opposing effects on cue-driven behavior depending upon the estrous cycle stage. In contrast, both estrus and non-estrus rats exposed to repeated prior stress exhibited an increase in conditioned responding relative to animals that underwent a single stress exposure. We further examined if the distal stress experience subsequently impacted extinction and the ability to learn a new cue-reward association. Prior stress did not affect extinction, though estrus and non-estrus rats exposed to repeated prior stress exhibited higher levels of conditioned responding to the novel cue-reward pairing. Taken together, our data demonstrate that the influence of stress on reward learning is impacted acutely by the estrous cycle as well as by one’s prior history with stress.

## Introduction

Stress produces transient physiological responses which can lead to long-lasting behavioral changes. These persistent behavioral alterations can arise from stress impacting learning processes. Across species, stress elicits greater activation of the hypothalamic-pituitary-adrenal (HPA) axis in females relative to males [1,2]. This increased stress responsivity is thought to contribute to the higher incidence of stress-related disorders in women compared to men [1-3]. While prior studies have characterized the effect of stress on learning cue-outcome associations in males, it is also critical to determine how stress affects learning in females [4-13]. The impact of stress on learning has traditionally been studied in the context of stress-enhanced fear/threat learning [4-10,14-25]. In female rodents, a single stress experience enhances Pavlovian conditioned responding to cues associated with a foot shock [20,22-24]. Multiple exposures to stress can further enhance fear/threat learning [26]. Robust behavioral responding to aversive cues may initially appear beneficial, as it helps avoid threats in one’s environment. However, this stress-enhanced responding is accompanied by negative consequences. For example, stressed females also exhibit deficits during extinction [8,15,17,19-21,27].

Considerably less is known about the impacts of stress on reward learning. In human studies that included women but did not separate data by gender, stress prior to conditioning results in enhanced reward learning compared to controls [28-32]. A study that examined women specifically found that individuals who exhibited the greatest stress response also displayed the highest degree of conditioned responding to a reward [32]. Furthermore, stress experience produces deficits in extinction to a reward-paired cue in humans [33,34]. Interestingly, others have found that stress has no impact on reward learning in human women or female rats [35-39]. These conflicting findings may be due to an interaction of cycling hormones and stress.

Hormonal cycles, such as the menstrual cycle in humans and the estrous cycle in rodents, can modulate reward learning [40-47]. Women in the late luteal phase of the menstrual cycle during training on a probabilistic learning task display greater reward conditioning relative to those in the late follicular phase [40]. In rats, females trained during estrus (the rodent analog of the late luteal phase) display greater cue-reward associations [41,42,48]. This enhanced association is evident a month after the initial conditioning session, demonstrating that estrous stage during the formation of cue-reward associations can have a long-lasting impact on behavior [42]. While the estrous cycle and stress separately influence reward learning, it is unclear how the two may interact to affect cue-reward associations.

To address this, we exposed naturally cycling female rats to either a single or repeated stress (or control procedures) prior to training on an appetitive Pavlovian conditioning task. During Pavlovian training sessions, rats learned to associate an audio cue with the delivery of a food reward. We subsequently altered the training parameters to assess how the distal stress experience affected the extinction of the previously established cue-reward relationship as well as the capacity to learn a novel cue-reward association. By monitoring the estrous cycle throughout training we determined how the stress interacted with the cycle stage to influence reward learning.

## Methods

### I. Subjects

The University of Texas at San Antonio Institutional Animal Care and Use Committee approved all procedures. Adult (60-70 days old) female Sprague Dawley rats (Charles Rivers and Inotiv) were pair-housed upon arrival, allowed *ad libitum* access to water and chow, and maintained on a 12 hr light/dark cycle. Prior to behavioral procedures, rats were either single- or pair-housed.

### II. Behavioral procedures

Rats were placed on mild dietary restriction to 90% of their free feeding weight, allowing for a weekly increase of 1.5%. Rats were regularly handled and estrous smears were obtained for one week prior to onset of behavioral experiments and after each Pavlovian conditioning session. All behavioral sessions occurred during the rats’ light cycle. Pavlovian conditioning occurred in operant boxes (MedAssociates) with a grid floor, a house light, a recessed food tray equipped with an infrared beam-break detector, and a white noise and tone generator. Rats underwent one habituation session in which they were placed in an operant box and received 20 un-signaled food pellets (45 mg, BioServ) delivered at a 90 s variable interval.

#### IIa. Restraint stress and control procedures

Rats were placed into four different treatment groups: single stress, single control, repeated stress, and repeated control. Prior to the first Pavlovian conditioning session, rats in the single stress/control groups underwent one restraint stress or control procedure. The stress procedure involved confining the rats to a clear acrylic tail vein restrainer (Braintree Scientific) for 20 min [13,49]. For the control procedure, rats were placed in a clean, empty cage for 20 min [13]. Our control procedure was designed to be as similar as possible to the stress procedure (animals are removed from home cage and placed in a clear plastic container), so that the only variable between the groups was whether the animal was restrained (stress) or not (control). Both control and stress procedures took place in an isolated room, distinct from where Pavlovian conditioning sessions would subsequently occur. To avoid potential confounds due to social transmission of stress, cagemates underwent the same stress/control procedures[50-54]. Additionally, stress and control treatments were administered on separate days. Repeated control/stress treatment animals underwent stress or control procedure once a day for five consecutive days. A subset of repeated control rats underwent a handle only control procedure. These rats were removed from their cage, weighed, and placed back in their home cage once a day for five days. However, we found no significant difference between our standard control procedure and the handle only control procedure (main effect F_(1,25)_ = 1.6, p = 0.2), so those rats are combined as the repeated control groups. The first Pavlovian conditioning session began 20 min after the conclusion of the last stress or control procedure.

#### IIb. Pavlovian conditioning and contingency change

After their final stress or control treatment, rats were transferred to a novel, clean recovery cage for 5 min. They were then placed in the operant chamber for 15 min prior to beginning Pavlovian conditioning, for a total of 20 min between stress/control procedure and onset of Pavlovian conditioning [13]. Prior research in male rats demonstrated that stress-enhanced conditioned responding was evident 20 min after stress, but absent 2 hrs after stress [13]. Importantly, stress or control procedures were only administered prior to the first session. Rats underwent an additional 9 Pavlovian training sessions, for a total of 10 Pavlovian conditioning sessions. Each Pavlovian training session consisted of 50 trials. In each trial, rats were presented with a 5 s white noise conditioned stimulus (CS+; 70 dB) which co-terminated with the delivery of a single food pellet unconditioned stimulus (US) and a 4.5 s illumination of the tray light. Trials were separated by a 55 ± 15 s intertrial interval (ITI).

A subset of rats underwent further conditioning sessions where the cue-reward contingencies were altered. In these training sessions, the white noise cue that had previously indicated a reward was no longer rewarded (CS+ → CS-; CS-trial), while a novel audio cue (tone, 5 s; 70 dB) now predicted a reward delivery (new CS+; CS+ trial). Trials were separated by a 55 ± 15 s ITI. Training sessions following the cue-reward contingency change consisted of 25 CS-trials and 25 CS+ trials presented in a pseudo-random order. Rats underwent five of these additional training sessions. Following the completion of Pavlovian training, rats were assessed for anxiety-like behavior and general locomotor activity. Rats were placed in an elevated zero maze apparatus for 5 min. Total distance travelled and percent time spent in open and closed arms were measured.

#### IIc. Behavioral analysis

We recorded head entries into the food tray across training sessions. Conditioned responding was quantified as the change in rate of head entries during the 5 s CS relative to the 5 ss preceding the CS [13,55-58]. Response latency was calculated as the time difference between CS onset and the first head entry during the CS. The rate of head entries when the cue was not playing (non-CS head entry rate) served as a proxy measurement of general motor activity within Pavlovian conditioning sessions. Conditioned responding to the CS-during the cue-reward contingency change sessions was normalized to the level of responding during the first contingency change session (conditioning session 11) to account for the differences in conditioned responding between treatment groups. For a direct measure of motor activity and anxiety-like behavior, we recorded the total distance travelled and percent time spent in the open arms of the zero maze apparatus, respectively.

### III. Vaginal cytology and Estrous Cycle Tracking

Vaginal cells were obtained via lavage, which entails flushing saline into the vaginal canal using a pipette. The saline with the vaginal sample was then placed on a slide, allowed to air dry, and then stained with crystal violet [59]. The resulting estrous smears were then examined via light microscopy. The person evaluating the samples was blind to the stress status of the rat. Samples with >80% nucleated epithelial cells were considered proestrus, >80% cornified epithelial cells were estrus, >80% leukocytes were diestrus, and an approximately equal mixture of all described cells were metestrus [48,59].

Estrous smears were obtained daily for one week prior to behavioral experiments and after each Pavlovian conditioning session. On days in which rats underwent Pavlovian conditioning, estrous smears were obtained 30-60 min after the completion of the training session to minimize any potential impact of the estrous smear collection on conditioned responding. Females were categorized based on the estrous stage on the first day of Pavlovian conditioning (**Supplemental Fig. 1**). Rats were further categorized as estrus or non-estrus, consistent with others [60-63].

### IV. Statistical analysis

Statistical analyses were performed using GraphPad Prism. A repeated measures ANOVA was used to analyze effects on behavioral measures. In instances where animals did not complete the full training paradigm (due to experimenter error or other technical issues), a mixed-effects model was applied. Significance was set to α=0.05 for all tests. All data are graphed as mean ± SEM. When a significant main effect or significant interaction effect was found, a post-hoc Tukey’s multiple comparisons test was applied to identify statistical differences at the individual session level. All statistical analyses are presented in **Supplementary Table 1**.

## Results

### Single stress exposure regulates conditioned responding in an estrous cycle-dependent manner

Stress and the estrous cycle independently affect cue-outcome associations, but how the two may interact to regulate reward learning remains unknown [8,15,20-22,41-44,47]. To address this, naturally cycling female rats received either a single stress exposure (20 min restraint) or control procedure (20 min in an empty cage) 20 min prior to their first Pavlovian conditioning session (**Fig. 1A**). During the Pavlovian training sessions, rats were presented with a 5 s audio cue which co-terminated with the delivery of a food pellet (**Fig. 1B**). Prior research in male rats utilizing the same stress and conditioning paradigms found that a single stress enhances conditioned responding [13]. However, we found no effect of stress on conditioned responding and latency when all females were analyzed together (Stress: n = 34; Control: n = 37) (**Fig. 1C,F**). Additionally, there was no effect of stress on the non-CS head entry rate, which is a proxy measure for general motor activity during the Pavlovian training session (**Fig. 1I**). Given that the estrous cycle can influence cue-reward associations, we next separated the data based on the estrous stage each rat was in during their first Pavlovian conditioning session (Stress Non-estrus: n = 23; Stress Estrus: n=11; Control Non-estrus: n = 21; Control Estrus: n = 16) [41-43]. We found that a single stress exposure enhanced conditioned responding in non-estrus females (**Fig. 1D**). There was also an interaction effect of stress and session on the latency to enter the food port and the non-CS head entry rate in non-estrus rats (**Fig. 1G,J**). In contrast, we observed the opposite effect when stress was administered to females in estrus, as this led to a persistent decrease in conditioned responding (**Fig. 1E**). These findings demonstrate that a single exposure to stress impacts cue-driven behavior in an estrous-dependent manner.

**Figure 1.**
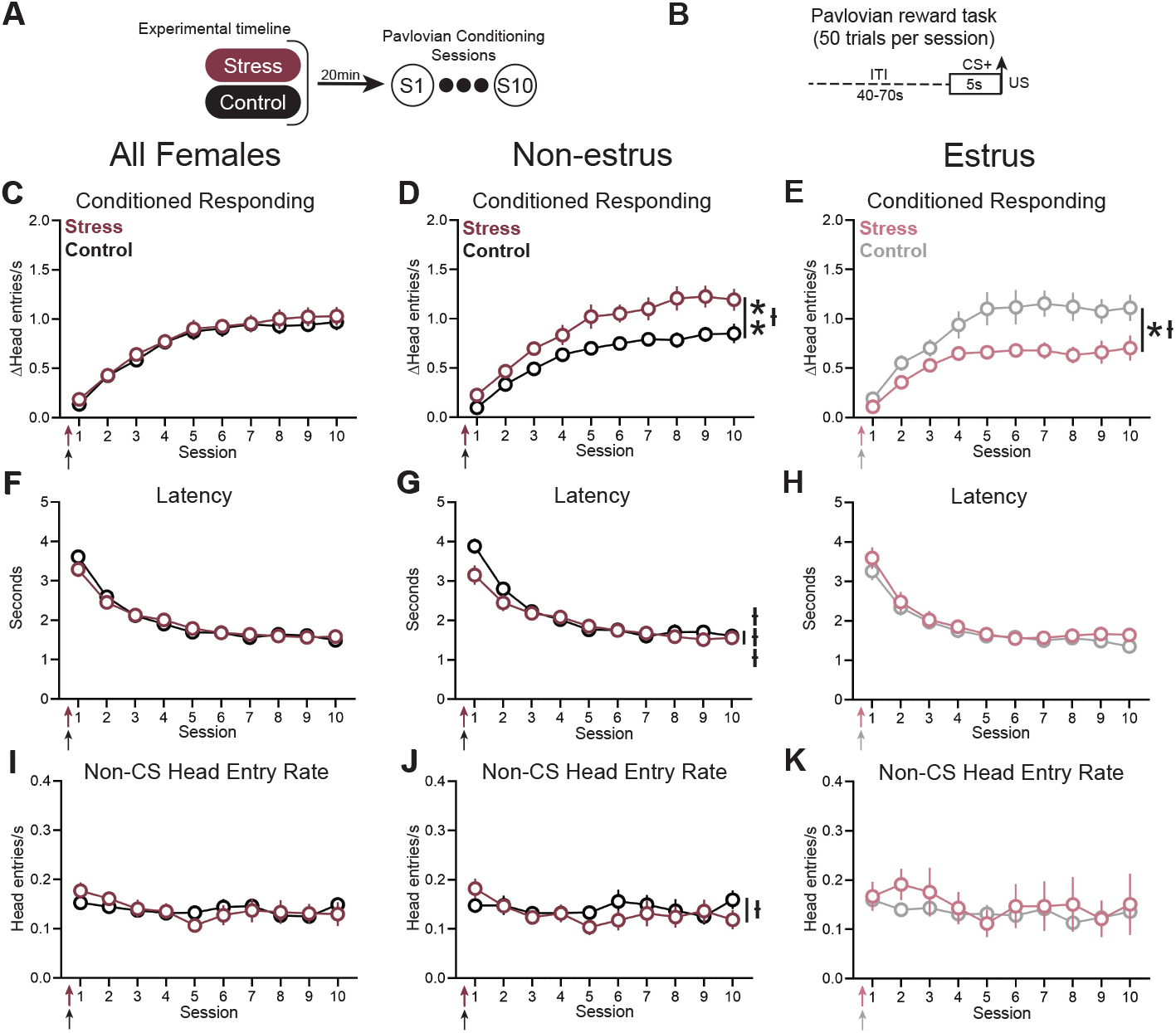
Estrous-dependent effect of a single stress exposure on conditioned responding. (A) Experimental timeline. Rats undergo a single restraint stress or control procedure prior to their first Pavlovian conditioning session. (B) Trial structure. Arrow indicates US delivery. (C) No effect of stress on conditioned responding when examining female rats across cycle stages (stress: n = 34; control: n = 37). (D) Stress enhanced conditioned responding in rats that underwent stress and the first Pavlovian conditioning session during a non-estrus phase of the estrous cycle (main effect of stress: F_(1,42)_ = 10.69, ** p = 0.002,; interaction of session x stress: F_(9,351)_ = 2.0, ƚ p = 0.04; non-estrus stress: n = 23; non-estrus control: n = 21). (E) Stress suppressed conditioned responding in rats that underwent their first Pavlovian conditioning session in estrus (main effect of stress: F_(1,25)_ = 5.9, * p = 0.02; interaction of session x stress: F_(9,218)_ = 2.021; ƚ p = 0.04; estrus stress: n = 11; estrus control: n = 16). (F) No effect of stress on response latency across all female rats. (G) Effect of stress on response latency in non-estrus rats (interaction of session x stress: F_(9,351)_ = 3.5, ƚƚƚ p = 0.0004). (H) No effect of stress on response latency in estrus rats. (I) Stress did not affect the non-CS head entry rate across all female rats. (J) Effect of stress on the non-CS head entry rate in non-estrus rats (interaction of session x stress: F_(9,351)_ = 2.1, ƚ p = 0.03). (K) No effect of stress on non-CS head entry rate in estrus rats. Arrows represent administration of stress, which occurred prior to session 1.

### Repeated stress further enhances conditioned responding across the estrous cycle

Repeated stress can further potentiate stress-enhanced fear learning, compared to a single stress [26]. Therefore, we next examined the effect of repeated restraint stress, or control procedures, on reward learning (Repeated Stress Non-estrus: n = 17; Repeated Stress Estrus: n = 12; Repeated Control Non-estrus: n = 20; Repeated Control Estrus: n = 7). In the repeated treatment groups, restraint stress (or control procedure) was administered daily for five consecutive days and Pavlovian conditioning began 20 min after the conclusion of the final stress/control exposure (**Fig. 2A**). When we did not separate our data according to the estrous cycle, we found no main effect of repeated stress (n = 29) compared to repeated control (n = 27) (**Fig. 2B,E,H**). However, there was an interaction effect of stress and session on conditioned responding (**Fig. 2B**). This result was driven by the non-estrus females, which also exhibited an interaction effect of stress and session on conditioned responding (**Fig. 2C**). Repeated stress additionally decreased the latency in estrus rats and increased non-CS head entry rate in non-estrus rats (**Fig. 2G,I**).

**Figure 2.**
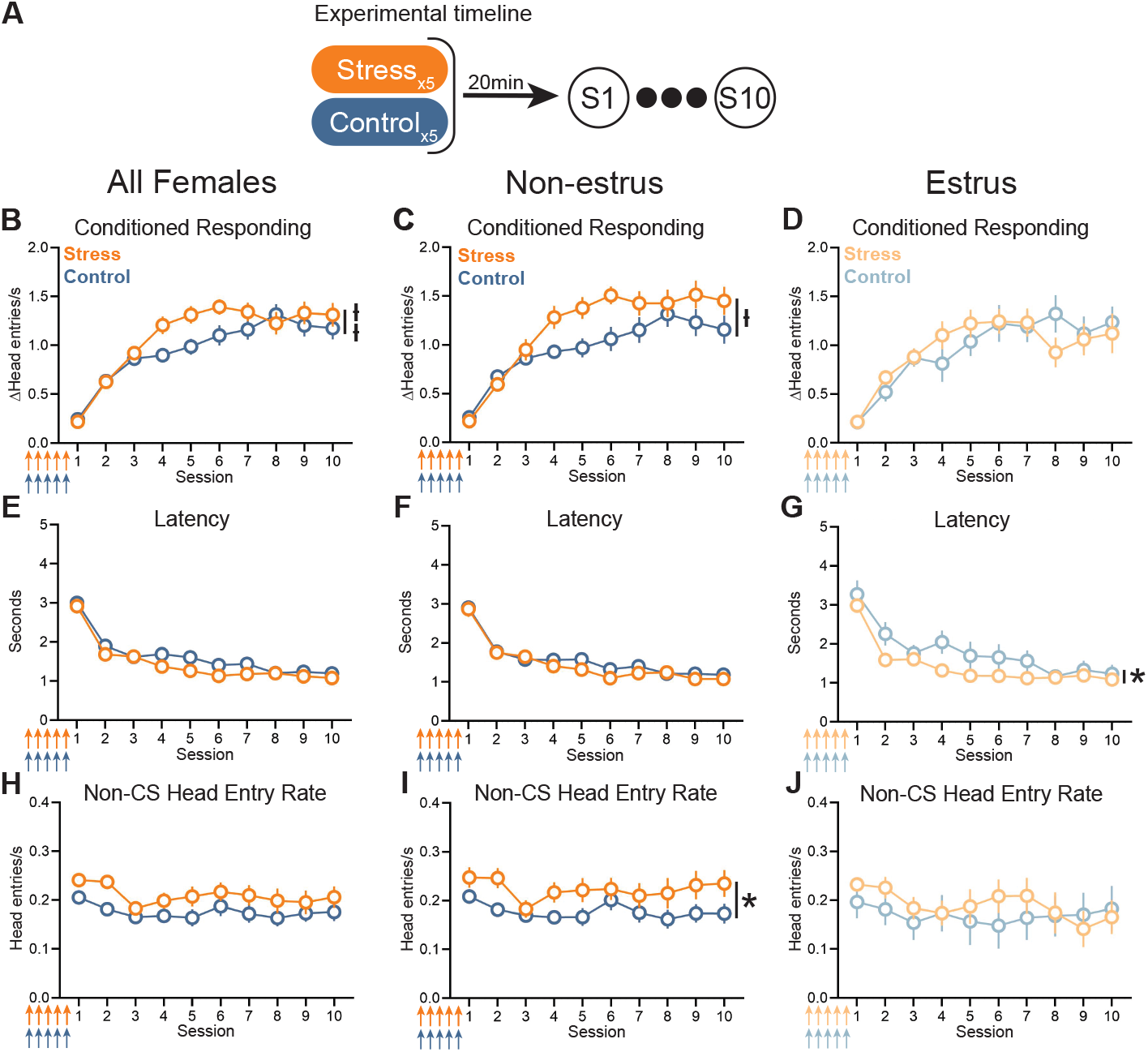
Repeated stress relative to repeated control treatment enhanced conditioned responding only in non-estrus rats. (A) Experimental timeline. (B) Effect of repeated stress on conditioned responding when examining female rats across cycle stages (interaction of session x stress: F_(9,464)_ = 2.8, ƚƚ p = 0.005; repeated stress: n = 29; repeated control: n = 27). (C) Repeated stress enhanced conditioned responding compared to repeated control treatment in non-estrus rats (interaction of session x stress: F_(9,302)_ = 2.5, ƚƚ p = 0.01; non-estrus repeated stress: n = 17; non-estrus repeated control: n = 20). (D) No effect of repeated stress relative to repeated control treatment on conditioned responding in estrus rats (estrus repeated stress: n = 12; estrus repeated control: n = 7). (E-G) Effect of repeated stress relative to repeated control treatment on response latency across all females (E), rats in non-estrus (F), and rats in estrus (G, main effect of stress: F_(1,17)_ = 4.7, * p = 0.04). (H-J) Effect of repeated stress relative to repeated control treatment on the non-CS head entry rate across all females (H), rats in non-estrus (I, main effect of stress: F_(1,35)_ = 4.3, * p = 0.04), and rats in estrus (J). Arrows represent administration of stress, which occurred prior to session 1.

We next compared the behavior of repeated stress/control animals to those that underwent a single stress/control treatment (**Fig. 3A; Supplemental Fig. 2A-B; Supplemental Fig. 3A**). Repeated stress resulted in a further enhancement of conditioned responding in both non-estrus and estrus rats (**Fig. 3B-C**). Focusing first on the non-estrus rats, we found that repeated stress decreased the latency to enter the food port and increased the non-CS head entry rate relative to rats exposed to a single stress (**Fig. 3D,F**). Interestingly, non-estrus rats that underwent repeated control treatments similarly exhibited enhanced conditioned responding and reduced latency relative to rats exposed to a single control treatment (**Supplemental Fig. 3B,D**). These data suggest that the repeated control treatment may function as a mild stressor in non-estrus rats. In estrus rats, repeated stress enhanced conditioned responding and reduced the response latency relative to estrus rats exposed to a single stress (**Fig. 3C,E**). However, there were no behavioral differences between estrus rats that underwent a single or repeated control treatment (**Supplemental Fig. 3C**,**E**,**G**). Taken together, our data suggest that repeated stress administration enhances reward associations, compared to a single stress, but also increases general activity in non-estrus rats.

**Figure 3.**
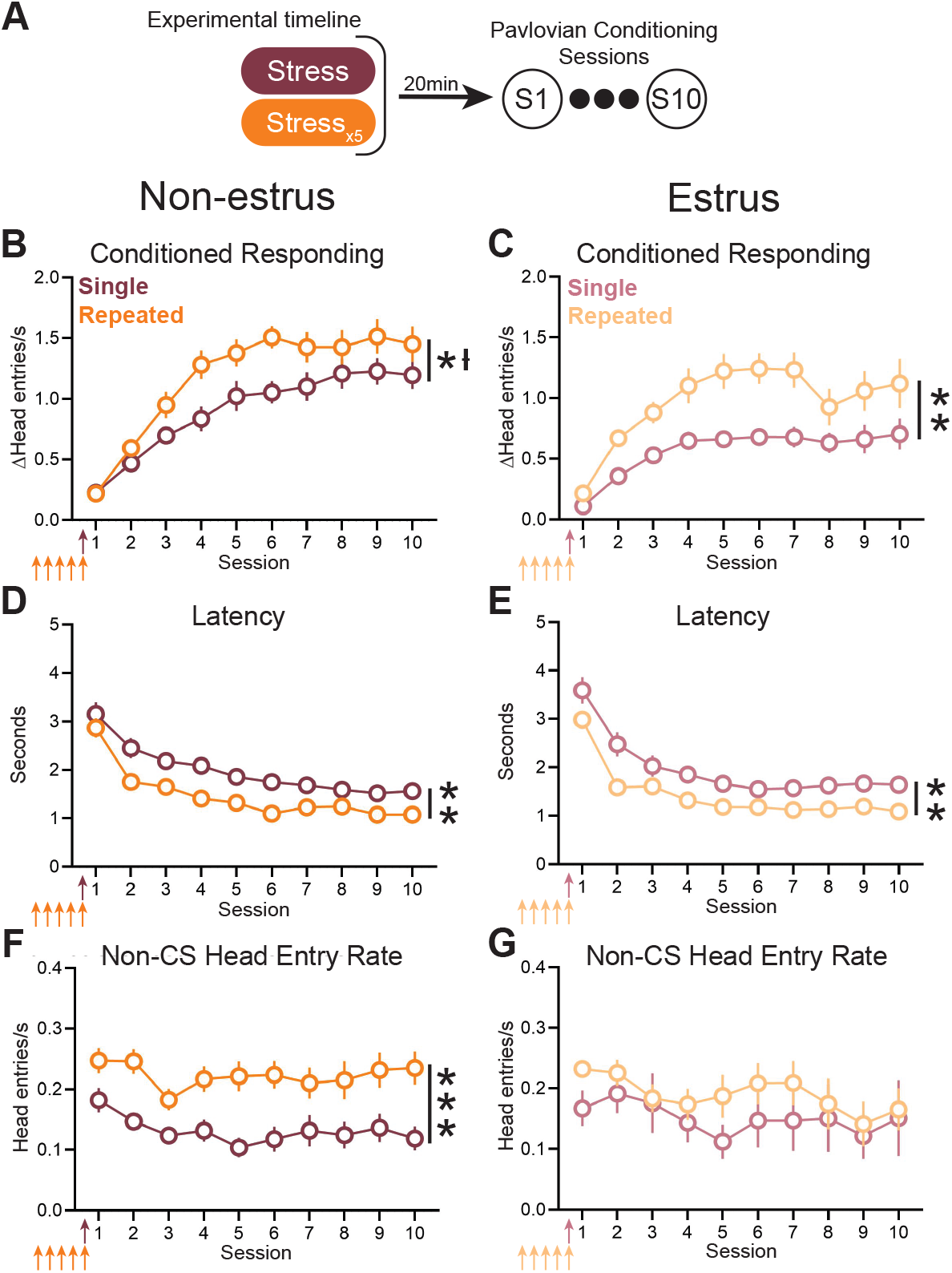
Repeated prior stress enhanced conditioned responding in estrus and non-estrus rats. (A) Experimental timeline. (B,C) Repeated stress enhanced conditioned responding relative to a single stress in non-estrus rats (main effect of repeated stress: F_(1,38)_ = 5.3, * p = 0.027, interaction of session and repeated stress: F_(9,308)_ = 2.20, ƚ p = 0.02; non-estrus single stress: n = 23; non-estrus repeated stress: n = 17) and estrus rats (main effect of repeated stress: F_(1,21)_ = 12.4, ** p = 0.002, Estrus single stress: n = 11; Estrus repeated stress: n = 12). (D,E) Repeated stress decreased the response latency in non-estrus rats (main effect of repeated stress: F_(1,38)_ = 8.4, ** p = 0.006) and estrus rats (main effect of repeated stress: F_(1,21)_ = 14.6, ** p = 0.001). (F,G) Repeated stress increased the non-CS head entry rate compared to single stress treatment only in non-estrus rats (main effect of repeated treatment: F_(1,38)_ = 16.5, *** p = 0.0002). Arrows represent administration of stress, which occurred prior to session 1.

The state of the animal when it is initially acquiring cue-outcome associations influences the resulting cue-driven behavior [13,43,64]. For example, when male rats are stressed 20 min prior to Pavlovian conditioning, they display long-lasting enhanced conditioned responding [13]. However, if stress is administered 2 hrs prior to conditioning, this enhancement is lost [13]. Accordingly, our initial analysis of rats undergoing repeated stress/control treatments focused on the estrous cycle stage on the last treatment day and first day of Pavlovian conditioning. However, we additionally examined whether the estrous cycle stage on the first day of the stress/control treatment influenced conditioned responding. As our repeated stress and control procedures lasted five days and the estrous cycle is a 4-5 day cycle, one might expect that many rats were in the same stage during their first and last day of stress or control procedure [48,59]. Indeed, 70% of repeated control rats and 66% of repeated stress rats were in the same estrus or non-estrus category on their first and last day of control/stress treatment. Accordingly, the behavioral findings are similar if the data are analyzed by estrous stage during the first or fifth day of stress/control treatment (**Supplemental Fig. 4**).

Previous research illustrates that stress can dysregulate the estrous cycle [65]. When analyzing the vaginal cytology of repeated stress/control rats, we observed a subset of rats would become “stuck” in a particular stage during the 5-day stress/control procedure, with 47.6% stuck in estrus and 52.4% in non-estrus (**Supplemental Fig. 5A**). We defined a stuck rat as one that repeated the same estrous stage for 3 or more consecutive days. The estrous cycle disruption in stuck rats most commonly began on the first day of stress/control treatment (day 1: 42.8%; day 2: 33.3%, day 3: 23.8%; **Supplemental Fig. 5B**). All rats that would become stuck did so by the third stress/control treatment (**Supplemental Fig. 5B**). Further, once a rat became stuck in a particular stage, it had an 80% chance of remaining in that stage until the stress/ control treatment ended. Interestingly, 62.1% of repeated stress rats became stuck while only 11.1% of repeated control rats became stuck (**Supplemental Fig. 5C**). When repeated stress/ control procedures concluded, rats resumed their typical estrous cycle. These data collectively highlight how exposure to repeated stress can transiently disrupt the estrous cycle.

### Repeated control/stress treatment abolishes estrous-dependent effects of stress

To further investigate the effect of the estrous cycle on cue-reward associations, we re-examined our data by comparing rats in estrus and non-estrus stages within each manipulation - single control, single stress, repeated control, and repeated stress (**Fig. 4**; **Supplemental Fig. 6**). In animals undergoing a single control treatment, estrus rats exhibited a higher level of conditioned responding relative to non-estrus rats, consistent with others (**Fig. 4A**) [41,42]. However, in animals undergoing a single stress treatment, non-estrus rats display greater conditioned responding relative to estrus rats **(Fig. 4B**). There was no effect of estrous stage on latency and non-CS head entry rate in single stress/control treatment rats, demonstrating that the estrous cycle selectively impacts conditioned responding (**Supplemental Fig. 6A**,**B**,**E**,**F**). In contrast to the rats that underwent a single stress/control treatment, those that received repeated stress/control treatments exhibited no difference in conditioned responding between estrus and non-estrus groups (**Fig. 4C-D**). Our findings demonstrate that the estrous cycle gates the effect of a single stress experience on conditioned responding, but this estrous-dependent effect is absent in rats that had received multiple prior exposures to stressful stimuli.

**Figure 4.**
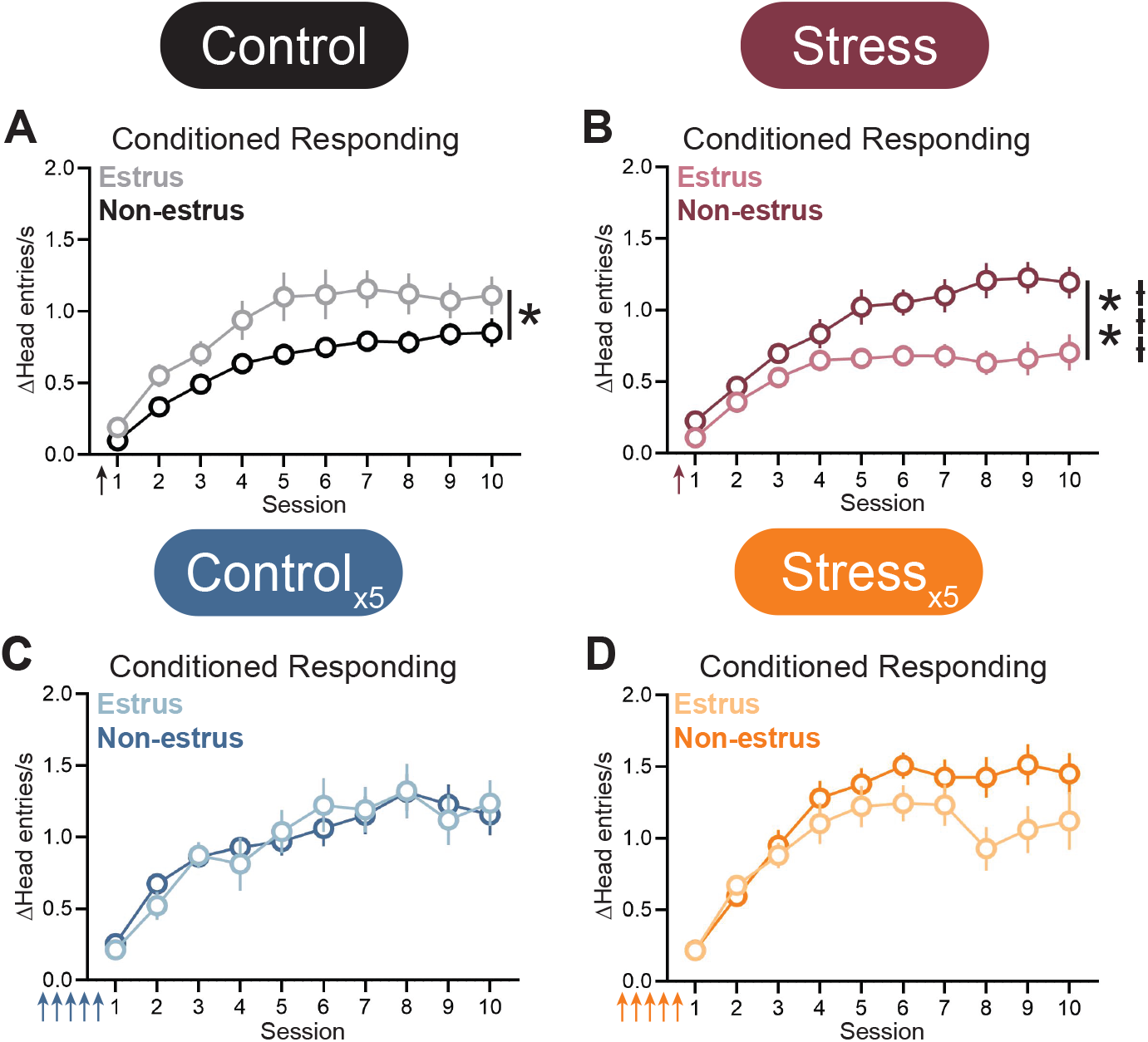
Effect of estrous stage on conditioned responding within manipulation groups. (A) In animals undergoing a single control treatment, estrus rats exhibited a higher level of conditioned responding relative to non-estrus rats (main effect of stage: F_(1,35)_ = 6.9, * p = 0.01). (B) In animals undergoing a single stress treatment, non-estrus rats exhibited a higher level of conditioned responding relative to estrus rats (main effect of stage: F_(1,32)_ = 9.2, ** p = 0.005; interaction of session x stage: F_(9,256)_ = 4.0, ƚƚƚ p < 0.0001). (C-D) No effect of estrous stage on conditioned responding in repeated control (C) or repeated stress rats (D). Arrows represent administration of stress, which occurred prior to session 1.

### Temporally distal repeated stress facilitates new reward learning without affecting extinction

Stress impairs extinction in fear conditioning tasks [8,11,17,19-21,27]. This suggests that prior stress exposure can have long-lasting impacts on flexible behavior temporally distant from when the stress was experienced. Here, we examined if stress experience prior to the first Pavlovian conditioning session would impact extinction as well as the ability to learn a new cue-reward association. After the initial 10 Pavlovian conditioning sessions in which the rats were trained to associate a white noise with a food reward, a subset of rats underwent a cue-reward contingency change. In the next five sessions, the previously rewarded white noise had no consequence (CS-) and a novel tone resulted in food delivery (CS+) (**Fig. 5A,F,K; Supplemental Fig. 7A**,**G**). We found no significant effect of stress, repeated treatment, or estrous stage on the extinction of conditioned responding to the CS-(**Fig. 5; Supplemental Figs. 7-9**). However, in rats that were in a non-estrus stage on the first day of Pavlovian conditioning, repeated stress enhanced conditioned responding to a new CS+ relative to both repeated control treatments (**Fig. 5I**) and a single stress (**Fig. 5N**). Similarly, repeated stress rats in estrus exhibited enhanced conditioned responding to a new CS+ relative to single stress estrus rats (**Fig. 5O)**. Many of the behavioral differences in the response latency to the CS+ and the non-CS head entry rate between treatment groups during the first 10 training sessions were evident in the training sessions after the contingency change (**Supplemental Fig. 10**). These long-lasting behavioral changes following repeated stress did not translate to alterations in anxiety-like behavior or general motor activity in a zero maze (**Supplemental Fig. 11**). Collectively, our results illustrate that repeated stress enhances conditioned responding soon after the stress exposures (**Fig. 2**,**3**) and subsequently facilitates the ability to acquire a new cue-reward association temporally distal to the stress exposures (**Fig. 5**).

**Figure 5.**
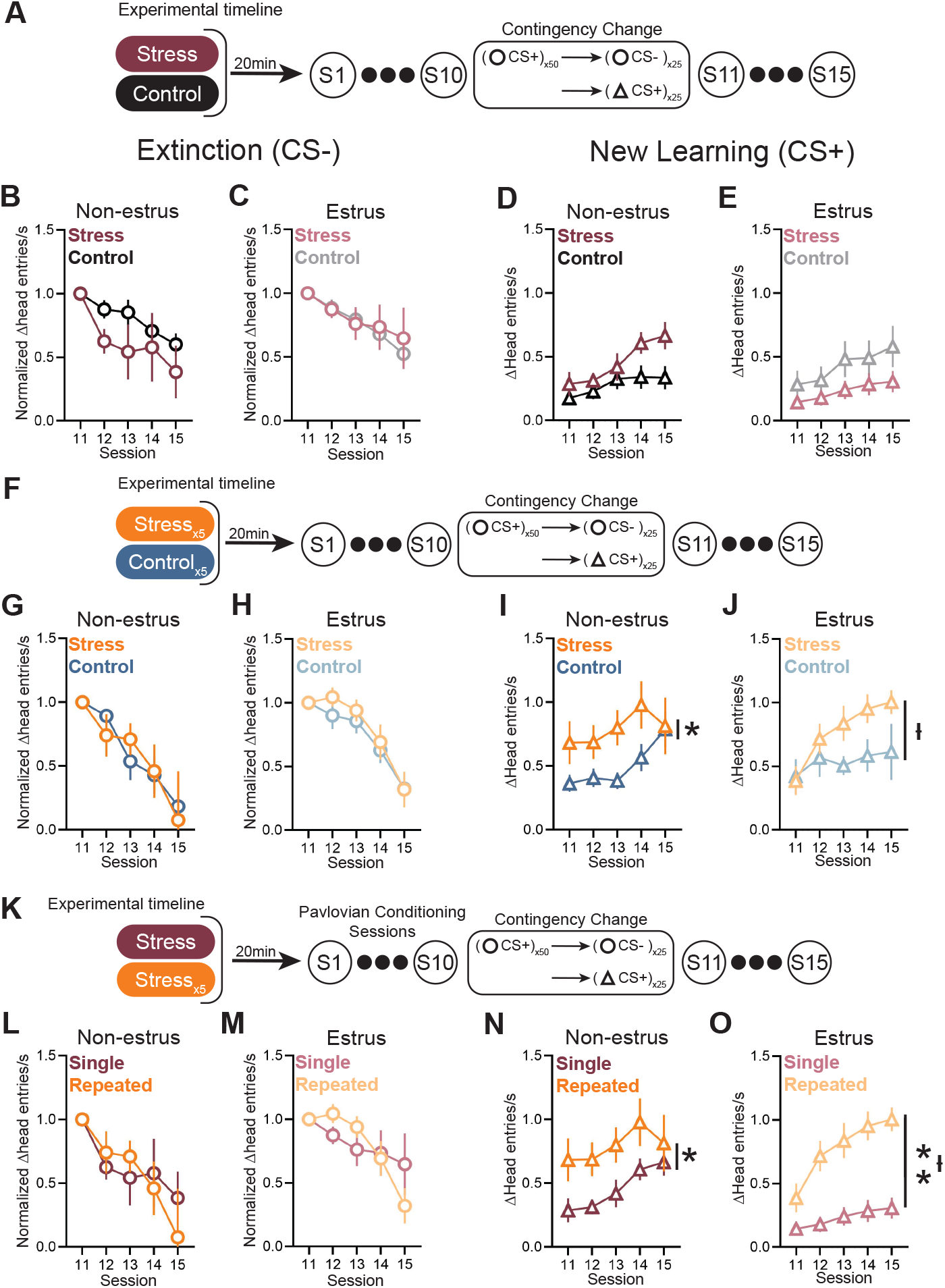
Temporally distal repeated stress facilitates new reward learning without affecting extinction. (A) Experimental timeline. (B-C) A single stress treatment (non-estrus: n = 6; estrus: n = 5) did not affect extinction relative to control treated rats (non-estrus: n = 8; estrus: n = 6). (D-E) A single stress treatment did not affect conditioned responding to a novel CS+ relative to control treated rats. (F) Experimental timeline. (G-H) Repeated stress treatment (non-estrus: n = 6; estrus: n = 10) did not affect extinction relative to the repeated control treatment (non-estrus: n = 8; estrus: n = 8) in either non-estrus (G) or estrus (H) rats. (I-J) Repeated stress enhanced conditioned responding to a new CS+ relative to the repeated control treatment in both non-estrus rats (I; main effect of repeated stress: F_(1,14)_ = 5.7, * p = 0.03) and estrus rats (J; interaction of session x repeated stress: F_(4,58)_ = 2.7, ƚ p = 0.04). (K) Experimental timeline. (L-M) Repeated stress treatment did not affect extinction relative to rats receiving a single stress treatment. (N-O) Repeated stress enhanced conditioned responding to a new CS+ relative to single stress in both non-estrus rats (N, main effect of repeated stress: F_(1,10)_ = 5.6, p = 0.04) and estrus rats (O, main effect of repeated stress: F_(1,13)_ = 11.9, ** p = 0.004; interaction of session x repeated stress: F_(4,50)_ = 3.1, ƚ p = 0.02).

## Discussion

Prior research demonstrates that a single stress exposure enhances conditioned responding to a reward-paired cue in male rats [13]. Here, we found that a single stress experience did not affect conditioned responding when examining female rats across all estrous cycle stages. However, this initial analysis obscured the opposing behavioral effect of stress on estrus and non-estrus rats. Specifically, female rats that began Pavlovian conditioning during non-estrus stages exhibited a stress-mediated *enhancement* in conditioned responding. In contrast, females rats that began Pavlovian conditioning during estrus exhibited a stress-mediated *suppression* in conditioned responding. Prior research performed in stress-naïve animals found higher levels of conditioned responding in estrus rats [42], which mirrors our findings from rats that underwent a single control treatment. Therefore, the effect of a single stress experience functionally inverts how the estrous cycle stage impacts cue-driven behavior in reward-based tasks.

Our data support previous literature demonstrating that the physiological state of an animal during learning influences future behavior. For example, the satiety and stress states of an animal during conditioning influence conditioned responding [13,66-68]. We propose that the impact of the estrous cycle stage on Pavlovian conditioned responding represents another example of state-dependent learning. While our interpretation of the data suggests that it is the state, whether estrous stage or stress status, of the animal during the first Pavlovian conditioning session that determines future conditioned responding, we do not see a difference in conditioned responding until later sessions. This is consistent with other studies that have found that differences in physiological state during early training can impact behavior in later sessions [13,67]. As such, our findings highlight the importance of the state of an animal during initial Pavlovian training. We propose that stress- and estrous-dependent alterations in cognition on the first day of training set the stage for differences in learning and memory consolidation, which manifest in later training sessions.

Repeated prior stress facilitates learning in fear/threat conditioning tasks [26]. Consistent with these findings, we found that repeated prior stress enhanced conditioned responding toward reward-predictive cues in both estrus and non-estrus rats when compared to rats receiving a single stress exposure. Furthermore, there was no difference in cue-driven behavior when directly comparing estrus and non-estrus rats that underwent the repeated stress treatment. These findings indicate that repeated stress experience blunts the influence of the estrous cycle on the behavioral effects of stress. We argue these results could explain why some human studies have not reported that cycle stage interacts with stress to modulate cue-based actions. A challenge with human research relative to preclinical studies is that it can be difficult to control for prior stress experience. Additionally, women and girls, compared to men and boys, consistently report higher daily stress levels, are more likely to experience stress, and are more likely to perceive an event as stressful [69-79]. Thus, human studies examining those with a menstrual cycle will likely include subjects with prior stress experience. Indeed, human studies that monitor the menstrual cycle often do not report an effect of cycle stage on stress-induced changes in cued associations [30,32,40,80-83]. Taken together, these findings highlight the importance of understanding when behavior is, and is not, gated by the estrous and menstrual cycles.

We found that non-estrus rats exhibit higher levels of conditioned responding following the repeated control treatment, relative to rats exposed to the single control treatment. Repeated stress (relative to a single stress) also increased conditioned responding in non-estrus rats. Furthermore, repeated stress produced higher levels of conditioned responding relative to repeated control treatments in non-estrus rats. These data collectively suggest that the repeated control treatment may function as a mild stressor in non-estrus rats. Alternatively, prior studies demonstrate that environmental enrichment can facilitate reward learning [84-86]. Therefore, the behavioral effect of the repeated control treatment may not be dependent upon stress per se, but rather may arise from experiencing the relatively novel control context. Future experiments will be needed to assess how physiological measures of stress, such as corticosterone levels, are related to single and repeated exposures to stress and control treatments. Importantly, there was no difference in the level of conditioned responding between single and repeated control treatments in estrus rats. These data further underscore the importance of examining how the estrous cycle stage during early learning can produce long-lasting behavioral consequences.

Based on the findings from stress-enhanced fear/threat learning models, one might expect that prior stress will alter behavioral responding during extinction [11,12,20]. In support, administering a cold pressor test prior to training on a reward-based task produced a transient deficit in performance during early extinction in humans [34]. However, neither a single nor repeated stress (compared to relevant control procedures) impacted extinction to the previously rewarded cue. Therefore, the impact of prior stress on extinction may depend on task-specific parameters such as the stressor used and whether the cue predicted an aversive or rewarding outcome. While stress did not affect extinction in our task, we instead found that prior repeated stress treatment enhanced conditioned responding (1) to the initial cue-reward pairing and (2) to a novel cue-reward pairing that was temporally distal to the stress exposure. This suggests that repeated prior stress experience coupled with reward learning primes rats to rapidly acquire a new cue-outcome association. It is possible that this enhanced learning is facilitated by the faster response latency or greater levels of unrewarded head entries displayed by the repeated stress rats during the initial ten Pavlovian training sessions.

While the data presented here provides valuable insight into the effects of stress and the estrous cycle on cue-reward associations, future studies are needed to identify the neural systems responsible for these behavioral outcomes. Acute stress activates the HPA axis, whereby promoting the release corticosteroids which elicit cellular responses throughout the periphery and the central nervous system. Repeated stress can dysregulate this canonical stress-response and is accompanied by stress-related neuroadaptations in a variety of brain regions including the prefrontal cortex, hippocampus, and ventral midbrain [87-89]. In particular, the mesolimbic dopamine system is well-positioned to mediate the interaction between stress, learning, and the estrous cycle. Dopamine release in the ventral striatum is necessary for Pavlovian conditioning to occur and dopamine is released in the ventral striatum as a result of restraint stress [90-93]. Furthermore, dopamine neuron activity fluctuates throughout the estrous cycle [41,61,94-97]. Additionally, we previously found that dopamine signaling in the ventral lateral striatum is necessary for stress-enhanced conditioned responding to a reward-paired cue in male rats [13]. Thus, dopamine signaling in the ventral striatum is well positioned to be the mechanism by which a single stress enacts estrous-dependent changes in reward conditioning in female rats as well.

Data from our lab and others demonstrate that exposure to stress promotes the expression of ‘goal-tracking’ conditioned responses in male rats [13,98]. Based on these findings one might predict that stress enhances the value of the rewarding outcome. However, stress can reduce sucrose preference, impair instrumental learning in reward-based tasks, and reduce the motivation to work for food rewards in male rats [49,99,100]. Therefore, the effect of stress on appetitive behaviors is task-specific. Our current results illustrate that sex and estrous cycle stage are also factors the determine how stress regulates behavior. Additionally, the behavioral outcome of stress is dependent upon the stressor type, stress intensity, and number of stress exposures [101-103]. Ultimately, the net effect of stress on behavior depends upon a complex interaction of the sex and hormonal status of the subject, the nature of the stressor, and the behavioral task utilized.

In conclusion, we found that a single stress elicited divergent effects on reward learning in an estrous-dependent manner. However, repeated stress enhanced conditioned responding, compared to a single stress, in an estrous-independent manner. Alterations in reward processing is a common symptom in many stress-related disorders, which disproportionately affect women [75,104-110]. As such, increasing research is needed to examine the impact of stress on cue-outcome associations in female subjects. The data presented here highlights the need for continued inclusion of female subjects and increased attention to the role of cycling hormones in pre-clinical studies investigating the effect of stress on behavioral outcomes.

## Acknowledgements

We would like to thank Claire Stelly for providing training with the behavioral task.

