## Supplemental Materials for "Estrous cycle stage gates the effect of stress on reward learning"

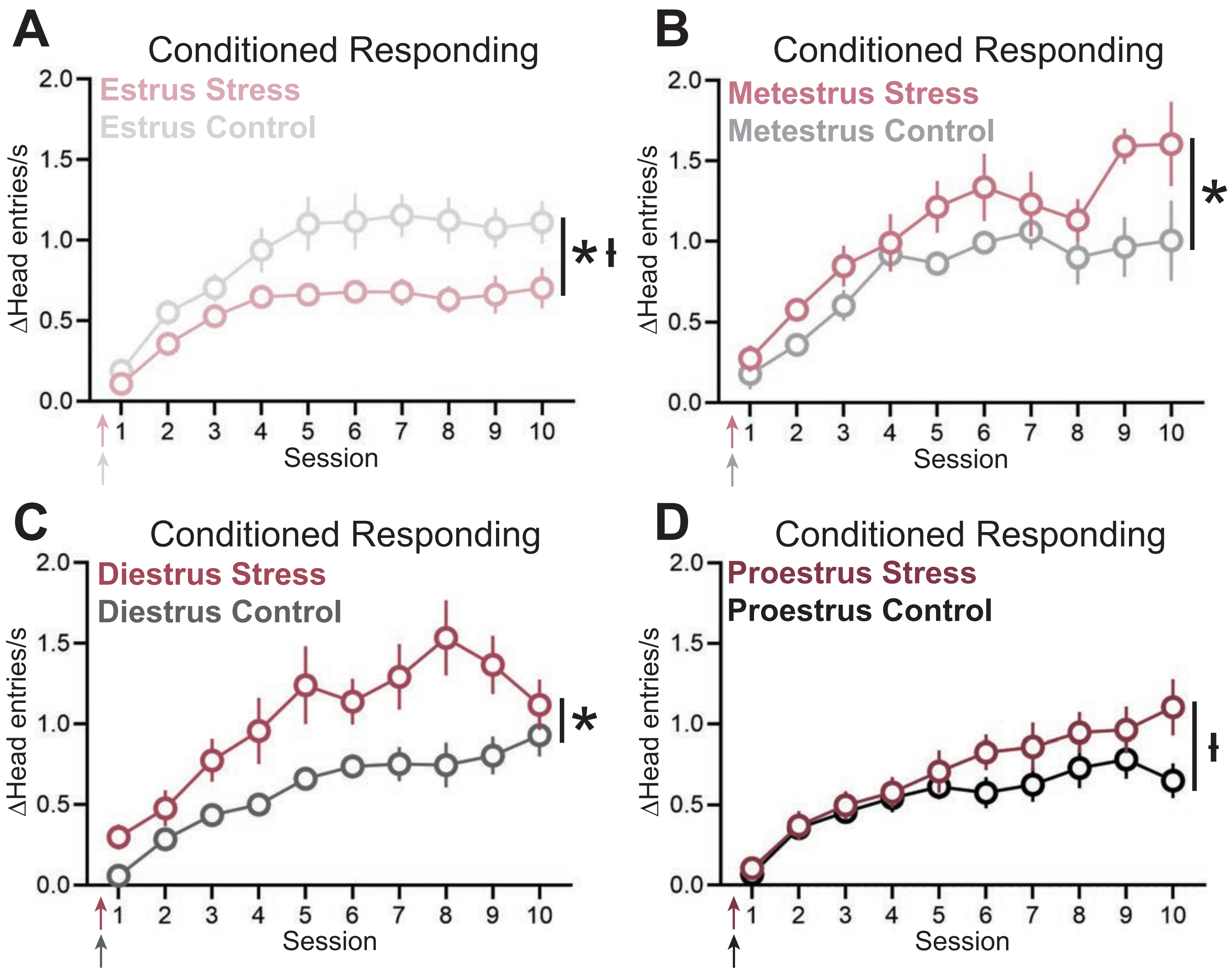


**Supplementary Figure 1:** Effect of a single stress or control treatment on conditioned responding during different phases of the estrous cycle. Estrous stage is based on stage during first Pavlovian conditioning session. (A) Stress suppressed conditioned responding in estrus rats (main effect of stress: $F_{(1,25)} = 5.9$, * $p = 0.02$; interaction of session x stress: $F_{(9,218)} = 2.0$, ł $p = 0.04$; control n = 16, stress n = 11) (B) Stress enhanced conditioned responding in metestrus rats (main effect of stress: $F_{(1,10)} = 7.3$, * $p = 0.02$; control n = 6; stress n = 6). (C) Stress enhanced conditioned responding in diestrus rats (main effect of stress: $F_{(1,14)} = 7.3$, $p = 0.02$; control n = 7, stress n = 9). (D) Effect of stress on conditioned responding in proestrus rats (interaction of session x stress: $F_{(9,125)} = 2.0$, ł $p = 0.04$; control n = 8; stress n=8). Arrows represent administration of stress, which occurred prior to session 1.

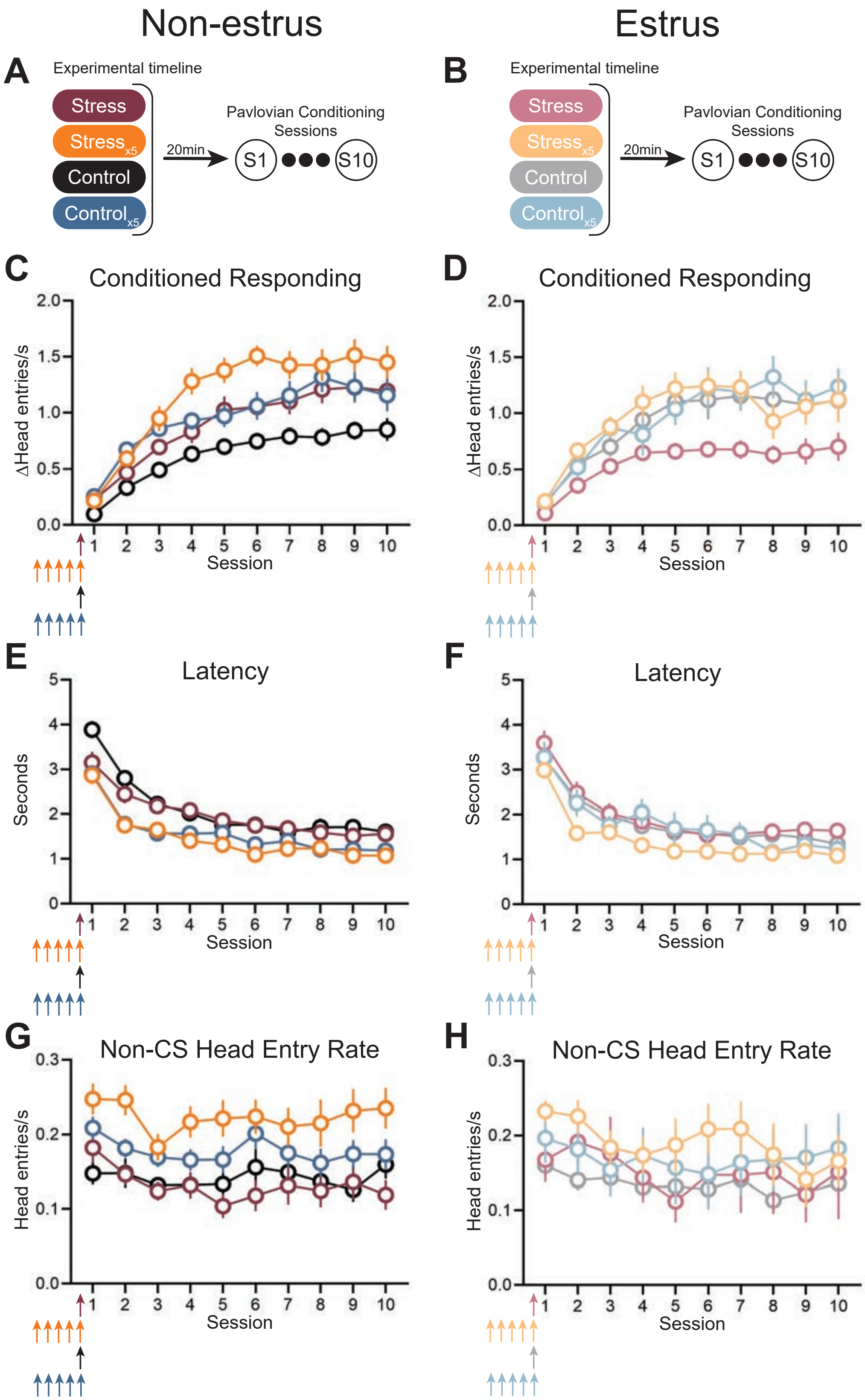


**Supplementary Figure 2:** Direct comparison of all four treatment groups in first ten sessions. (A) Experimental timeline for non-estrus animals. (B) Experimental timeline for estrus animals. (C) Conditioned responding of animals that began Pavlovian conditioning during non-estrus stages in single control, single stress, repeated control, and repeated stress conditions. A three-way mixed-effects model revealed a significant effect of repeated treatment ($F_{(1,77)}$ = 18.89, $p < 0.0001$) and stress treatment ($F_{(1,77)}$ = 12.68, p = 0.0006). Additionally, there was an interaction of session and stress ($F_{(9,653)} = 3.466$, $p = 0.0003$) and a three-way interaction of session, repeated treatment, and stress ($F_{(9,653)} = 1.399$, $p = 0.1846$). (D) Conditioned responding of animals that began Pavlovian conditioning during estrus in single control single stress, repeated control, and repeated stress conditions. A three-way mixed-effects model found a significant effect of repeated treatment ($F_{(1,42)} = 4.161$, $p = 0.0477$) and an interaction of session and stress treatment ($F_{(9,362)} = 1.981$, $p = 0.0405$). (E) Latency of non-estrus animals. A three-way mixed-effects model found a significant effect of repeated treatment ($F_{(1,77)} = 19.82$, $p < 0.0001$). There was also an interaction of session and repeated treatment ($F_{(9,653)} = 2.542$, $p = 0.0071$) and a three-way interaction of session, repeated treatment, and stress treatment ($F_{(9,653)} = 2.640$, $p = 0.00522$). (F) Latency of estrus animals. (G) Non-CS head entry rate of non-estrus animals. A three-way mixed effects model found a significant effect of repeated treatment ($F_{(1,77)} = 17.13$, $p < 0.0001$). (H) Non-CS head entry rate of estrus animals. Arrows represent administration of stress, which occurred prior to session 1.

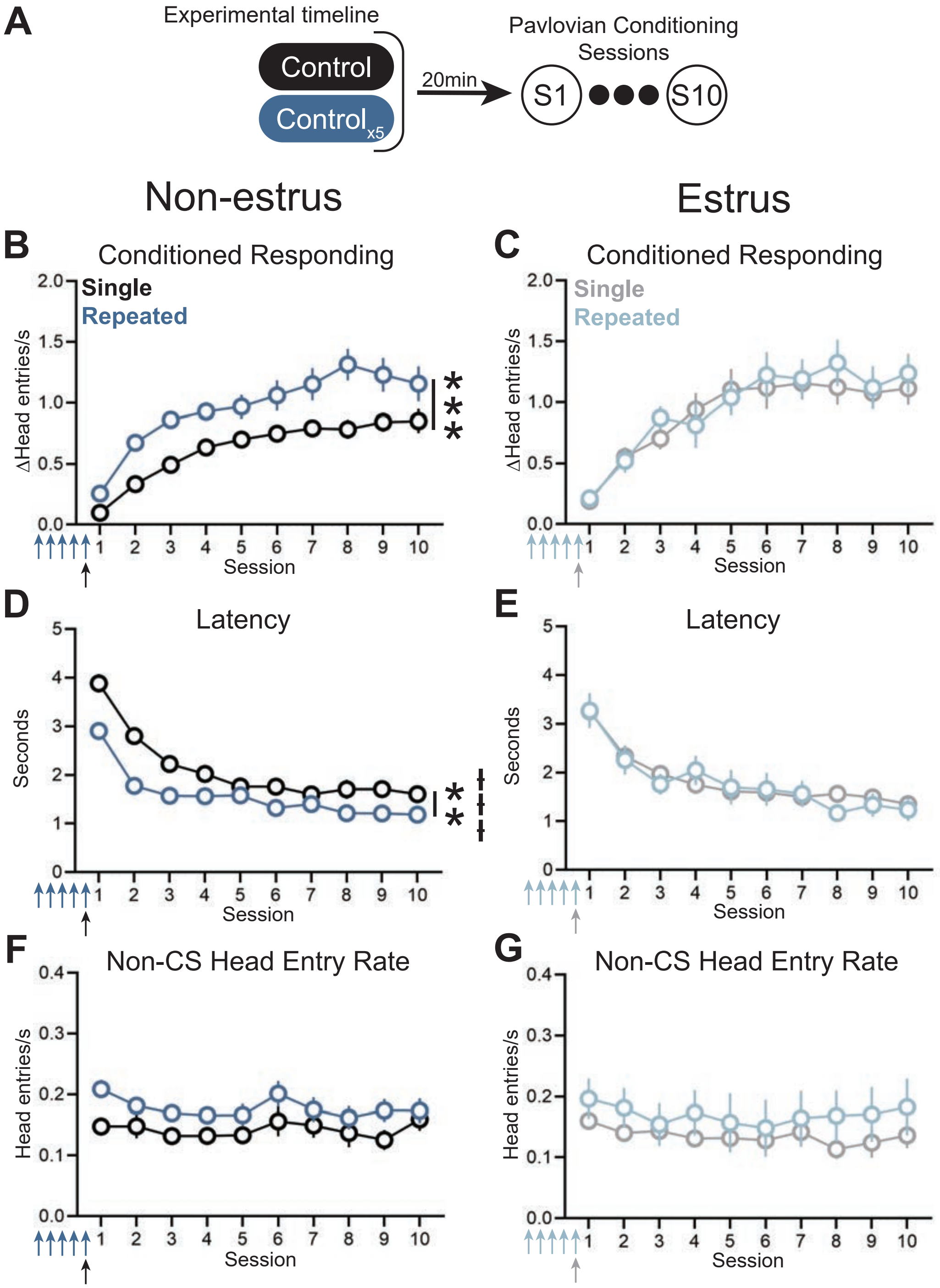


**Supplementary Figure 3:** Behavioral effects of single and repeated control treatments. (A) Experimental timeline. (B) Repeated control treatment in non-estrus rats enhanced conditioned responding relative to single control treatment rats (main effect of repeated treatment: $F_{(1,39)}$ = 17.3, *** p = 0.0002; single control: n = 21; repeated control: n = 20). (C) No difference in conditioned responding between single and repeated control estrus rats (single control: n = 16; repeated control: n = 7). (D) Repeated control treatment in non-estrus rats decreased the response latency relative to single control treatment rats (main effect of repeated treatment: $F_{(1,39)}$ = 11.8, p = 0.001; interaction of session and repeated treatment: $F_{(9,345)}$ = 4.3, ⧻ p <0.0001). (E) No difference in response latency between single and repeated control estrus rats. (F-G) No difference in the non-CS head entry rate latency between single and repeated control non-estrus rats (F) and estrus rats (G). Arrows represent administration of stress, which occurred prior to session 1.

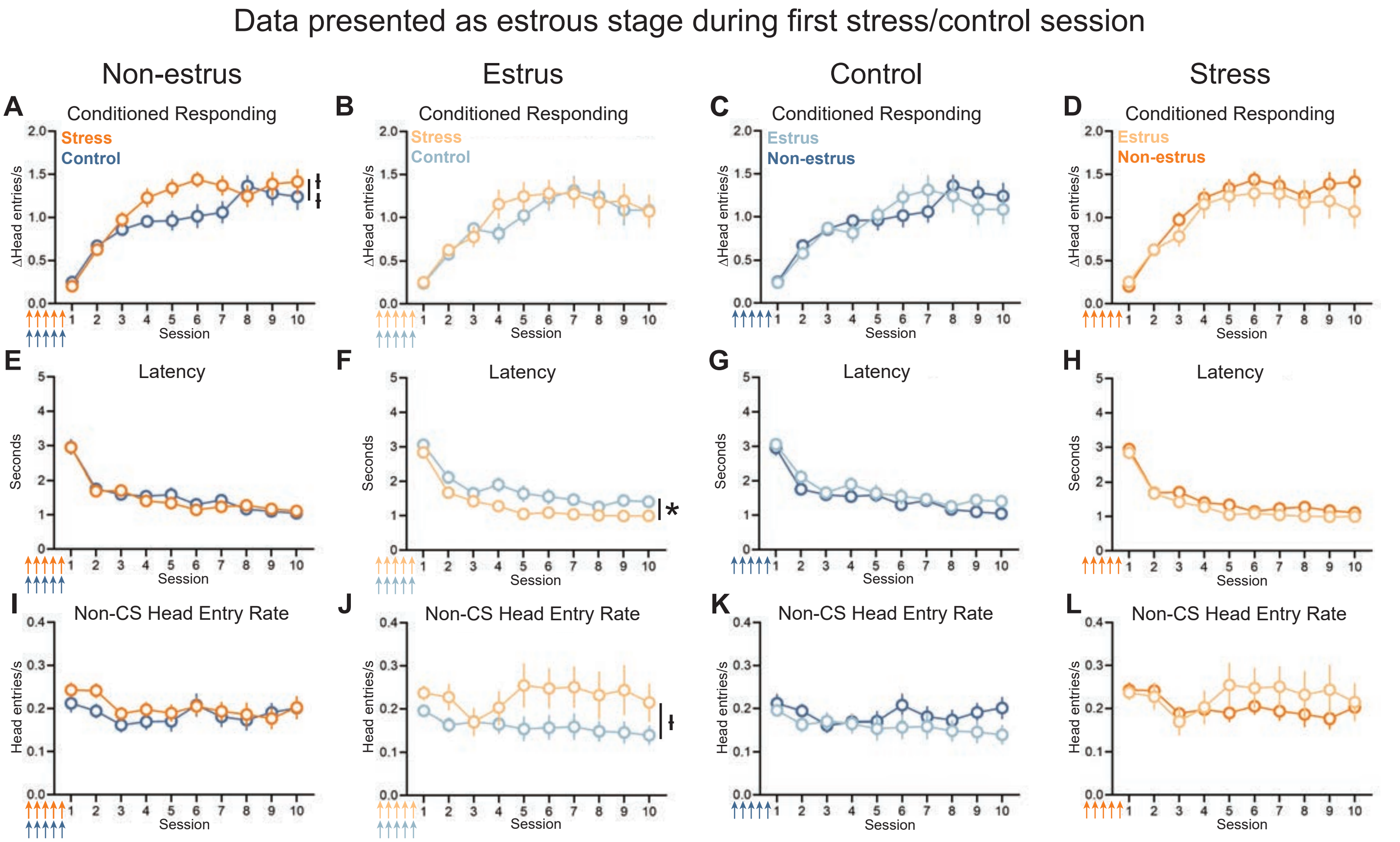


**Supplementary Figure 4:** Repeated treatment data analyzed by the estrous stage on the first stress/control treatment day. (A) Effect of repeated stress relative to repeated control treatment on conditioned responding in non-estrus rats (interaction of session x stress: $F_{(9,299)} = 2.8$, łł $p = 0.004$; control n = 16, stress n = 21). (B) No effect of repeated stress relative to repeated control treatment on conditioned responding in estrus rats (control n = 11, stress n = 8). (C-D) No effect of estrous stage on conditioned responding in repeated control (C) or repeated stress rats (D). (E) No effect of repeated stress relative to repeated control on responses latency in non-estrus rats. (F) Effect of repeated stress relative to repeated control on responses latency in estrus rats (main effect of stress: $F_{(1,17)} = 4.5$, * $p = 0.04$). (G-H) No effect of estrous stage on response latency repeated control (G) or repeated stress rats (H). (I) No effect of repeated stress relative to repeated control on the non-CS head entry rate in non-estrus rats. (J) Effect of repeated stress relative to repeated control on the non-CS head entry rate in estrus rats (interaction of session x repeated stress: $F_{(9,147)} = 2.1$, ł $p = 0.03$). (K-L) No effect of estrous stage on the non-CS head entry rate in control (K) or repeated stress rats (L). Arrows represent administration of stress, which occurred prior to session 1.

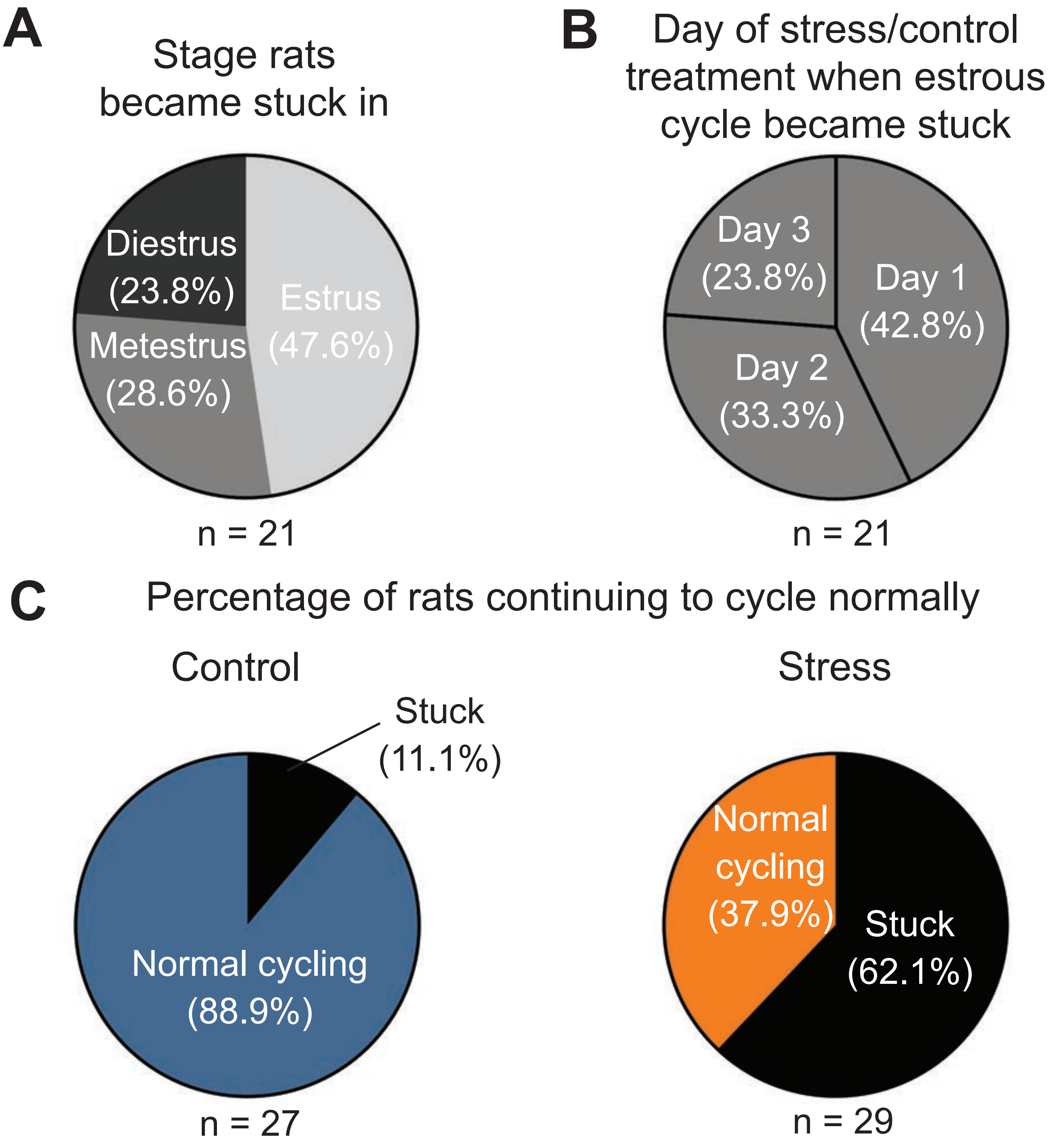


**Supplementary Figure 5:** Repeated stress/control treatment dysregulates the estrous cycle. Rats were classified as stuck if they remained in the same phase for 3+ days. (A) Percent of rats that became stuck in a given phase. (B) Day of stress or control treatment in which rats became stuck. 42.8% of rats became stuck in the phase they were in on the first day of stress/control treatment, 33.3% became stuck in their phase on the second day of treatment, and 23.8% became stuck in their phase on the third day of treatment. (C) Percentage of rats that continued to cycle normally or became stuck in a given phase during repeated control (left) or stress (right) treatment.

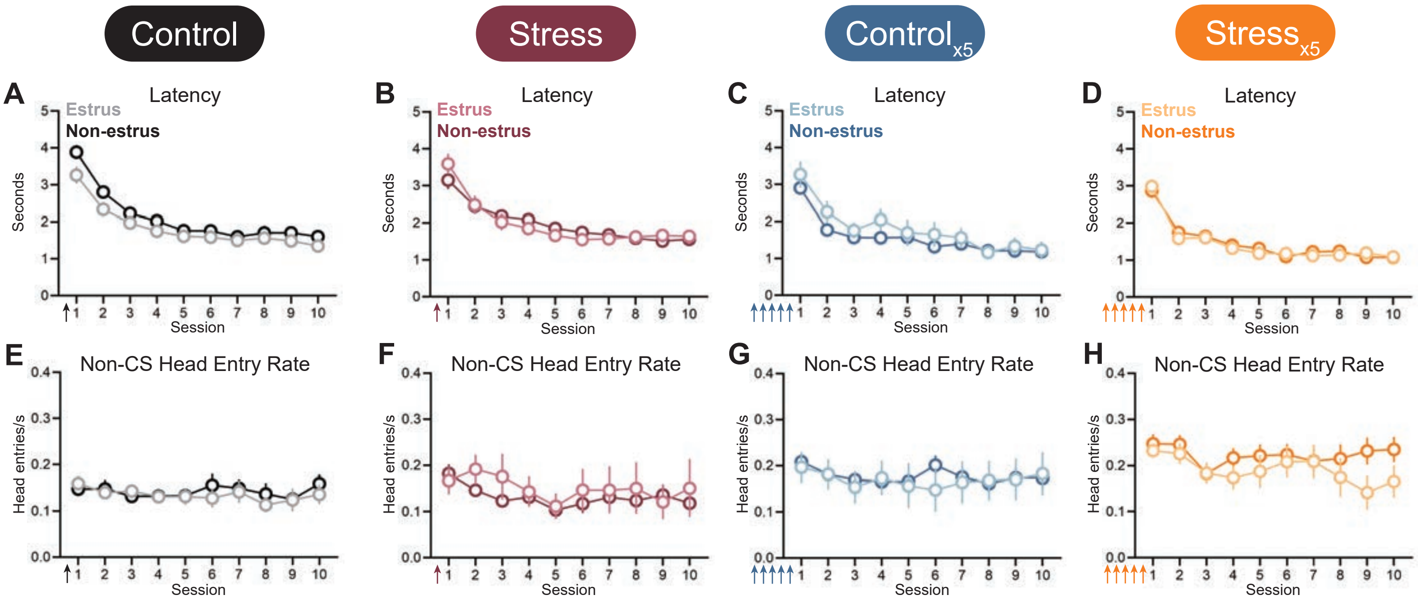


**Supplementary Figure 6:** Latency and non-CS head entry rate is not impacted by estrous cycle stage. (A-D) Estrous stage during first Pavlovian conditioning session did not impact the response latency for single control (A), single stress (B), repeated control (C), or repeated stress treatments (D). (E-H) Estrous stage during first Pavlovian conditioning session did not impact the non-CS head entry rate for single control (E), single stress (F), repeated control (G), or repeated stress treatments (H). Arrows represent administration of stress, which occurred prior to session 1.

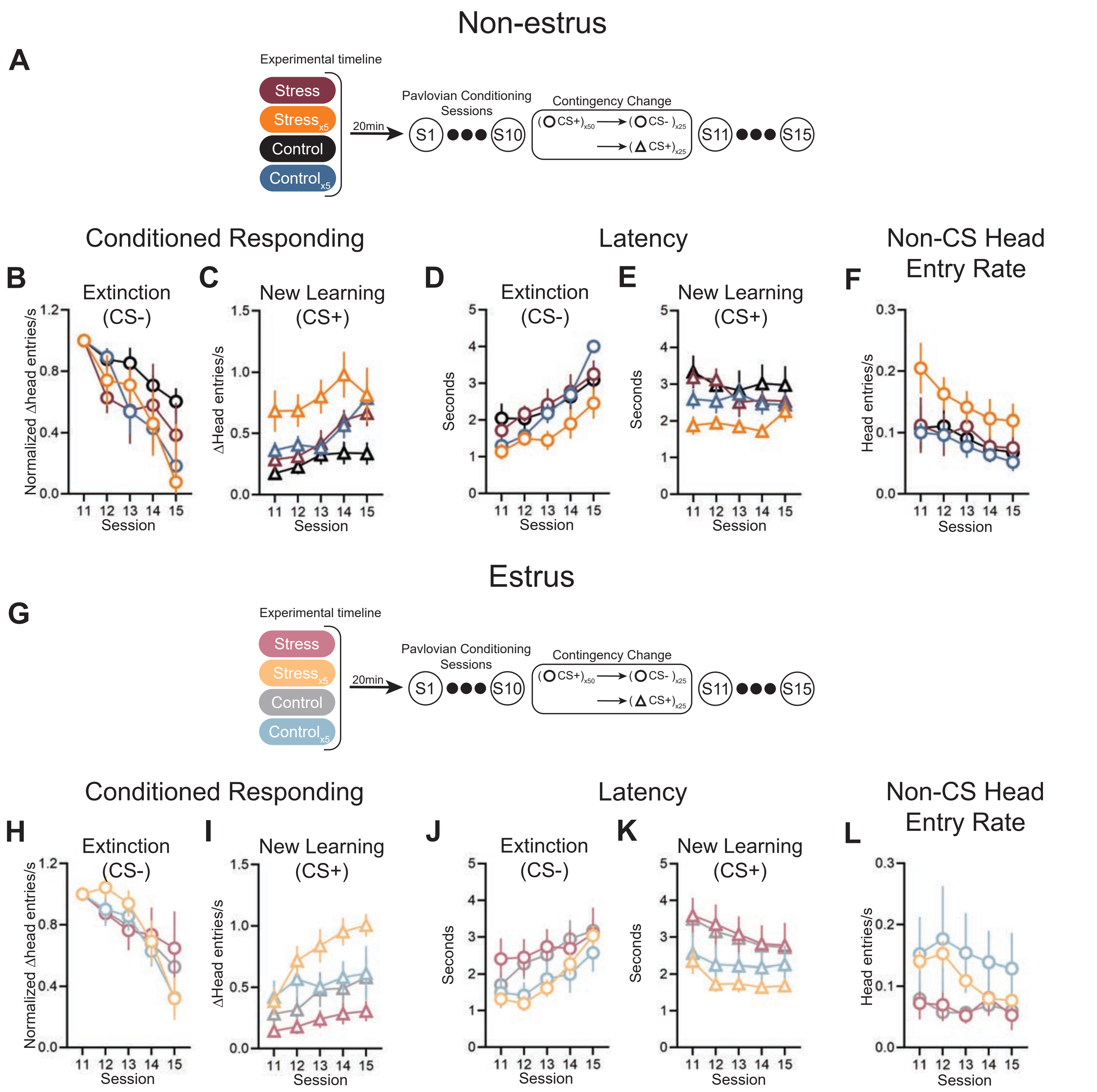


**Supplementary Figure 7:** Direct comparison of all four treatment groups after contingency change. (A) Experimental timeline for non-estrus animals. (B) Extinction, measured as conditioned responding to previously rewarded white noise, of non-estrus rats. A three-way mixed effects model found an interaction of session and repeated treatment ($F_{(4,95)}$ = 2.474, p = 0.0495). (C) New learning, measured as conditioned responding to a tone, of non-estrus rats. A three-way mixed effects model found a significant effect of repeated treatment ($F_{(1,26)}$ = 12.02, p = 0.0018) and of stress treatment ($F_{(1,26)}$ = 8.958, p = 0.0060). (D) Latency of non-estrus rats to enter the food port after the onset of the extinction cue. There was a three-way interaction of session, repeated treatment, and stress treatment ($F_{(4,95)}$ = 2.556, p = 0.0438). (E) Latency of non-estrus rats to enter the food port after the onset of the new cue. A three-way mixed effects model found a significant effect of repeated treatment ($F_{(1,26)}$ = 4.359, p = 0.0468). (F) Non-CS head entry rate of non-estrus rats did not significantly differ among repeated treatment or stress treatment groups. (G) Experimental timeline for estrus rats. (H) Extinction of estrus rats. A three-way mixed effects model found a significant interaction of session and repeated treatment ($F_{(4,93)} = 3.955$, $p = 0.0052$). (I) New learning of estrus rats. A three-way mixed effects model found a significant effect of repeated treatment ($F_{(1,24)} = 8.092$, $p = 0.0089$). (J) Latency of estrus rats to enter the food port after the onset of the extinction cue. A three-way mixed effects model found a significant effect of repeated treatment ($F_{(1,24)} = 4.507$, $p = 0.0443$) and a three-way interaction of session, repeated treatment, and stress treatment ($F_{(4,93)} = 2.620$, $p = 0.0398$). (K) Latency of estrus rats to enter the food port after the onset of the new cue. A three-way mixed effects model found a significant effect of repeated treatment ($F_{(1,24)} = 7.812$, $p = 0.0100$). (L) Non-CS head entry rate of estrus rats. A three-way mixed effects model found a significant interaction of session and repeated treatment ($F_{(4,93)} = 4.602$, $p = 0.0020$).

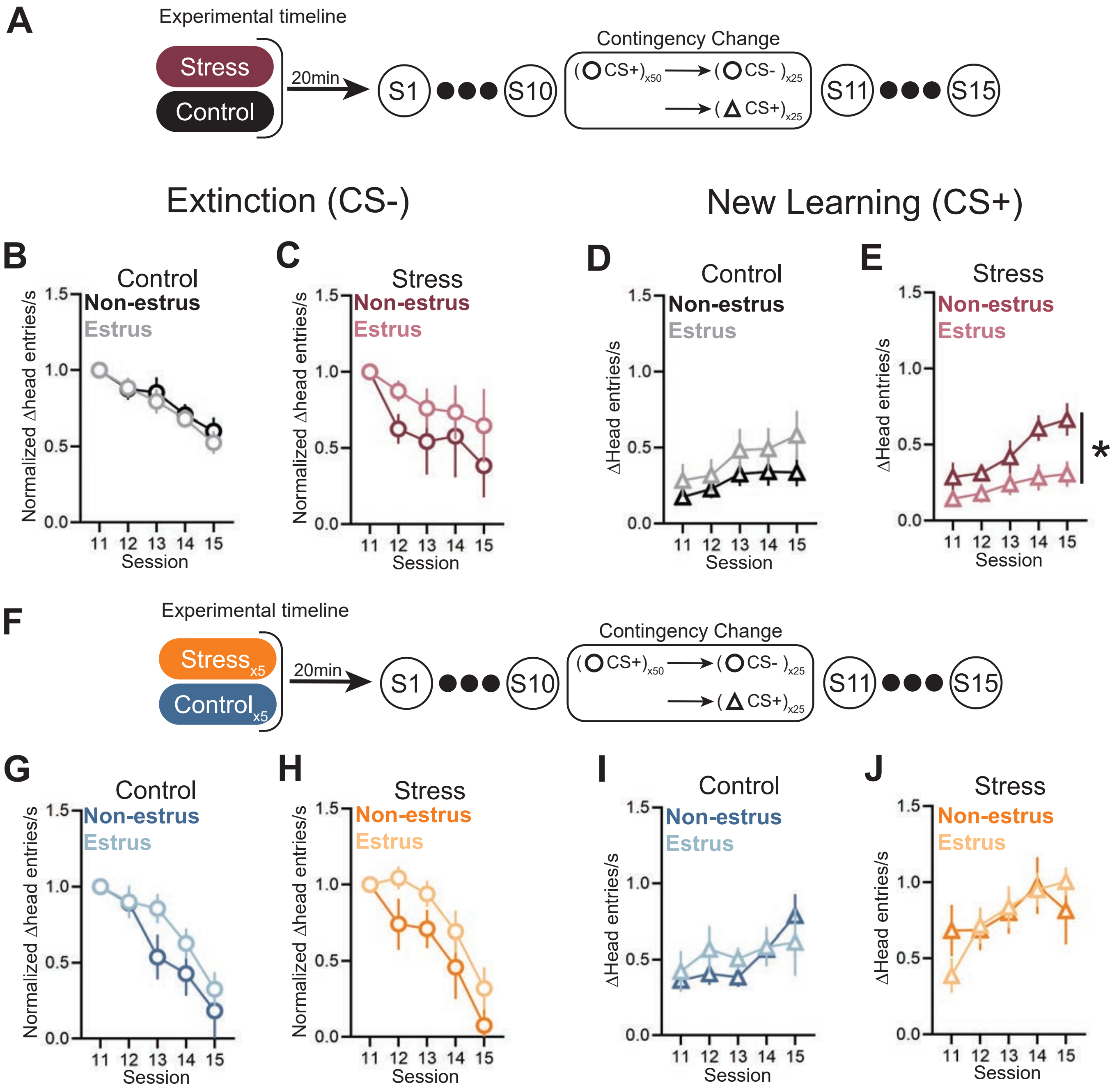


**Supplementary Figure 8:** Effect of estrous cycle stage during first Pavlovian conditioning session on extinction and acquiring a new cue-reward contingency. (A) Experimental timeline. (B-C) No effect of estrous cycle stage on extinction in single stress (non-estrus: n = 6; estrus: n = 5) (B) and single control (non-estrus: n = 8; estrus: n = 6) rats (C). (D) No effect of estrous cycle stage on conditioned responding to a new CS+ in single control rats. (E) Non-estrus rats that underwent a single stress prior to the first Pavlovian conditioning session exhibited enhanced conditioned responding to the new CS+ relative to estrus rats (main effect of stage: $F_{(1,9)} = 5.9$, * $p=0.04$). (F) Experimental timeline. (G-H) No effect of estrous cycle stage on extinction in repeated stress (non-estrus: n = 6; estrus: n = 10) (G) and repeated control (non-estrus: n = 10: estrus: n = 8) rats (H). (I-J) No effect of estrous cycle stage on conditioned responding to a new CS+ in repeated control (I) and repeated stress rats (J).

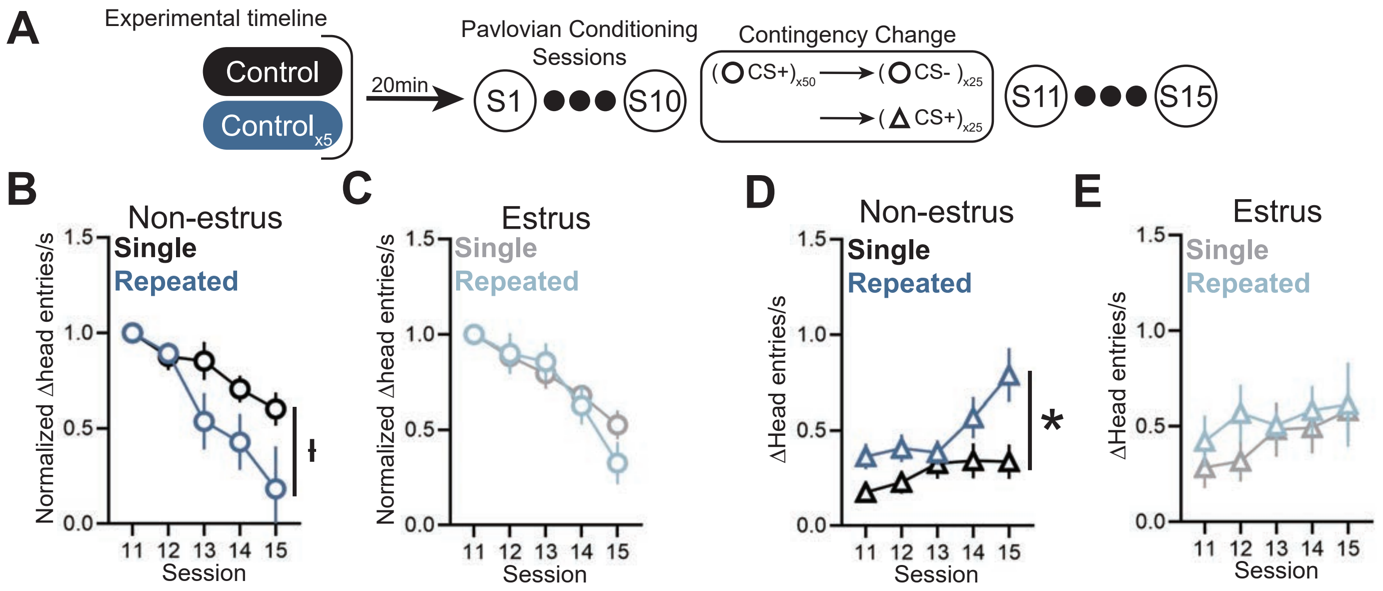


**Supplementary Figure 9:** Effect of repeated control compared to single control treatment on extinction and acquiring a new cue-reward contingency. (A) Experimental timeline. (B) Effect of repeated control relative to single control treatment on extinction in non-estrus rats (interaction of session x repeated: control treatment $F_{(4,58)}$ = 3.3, ł p = 0.02). (C) No effect of repeated control treatment relative to single control treatment on extinction in estrus rats. (D) Repeated control treatment enhanced conditioned responding to the new CS+ relative to single control treatment in non-estrus rats (main effect of repeated control treatment: $F_{(1,16)}$ = 5.6, * p = 0.03). (E) No effect of repeated control treatment on conditioned responding to the new CS+ relative to single control treatment in estrus rats.

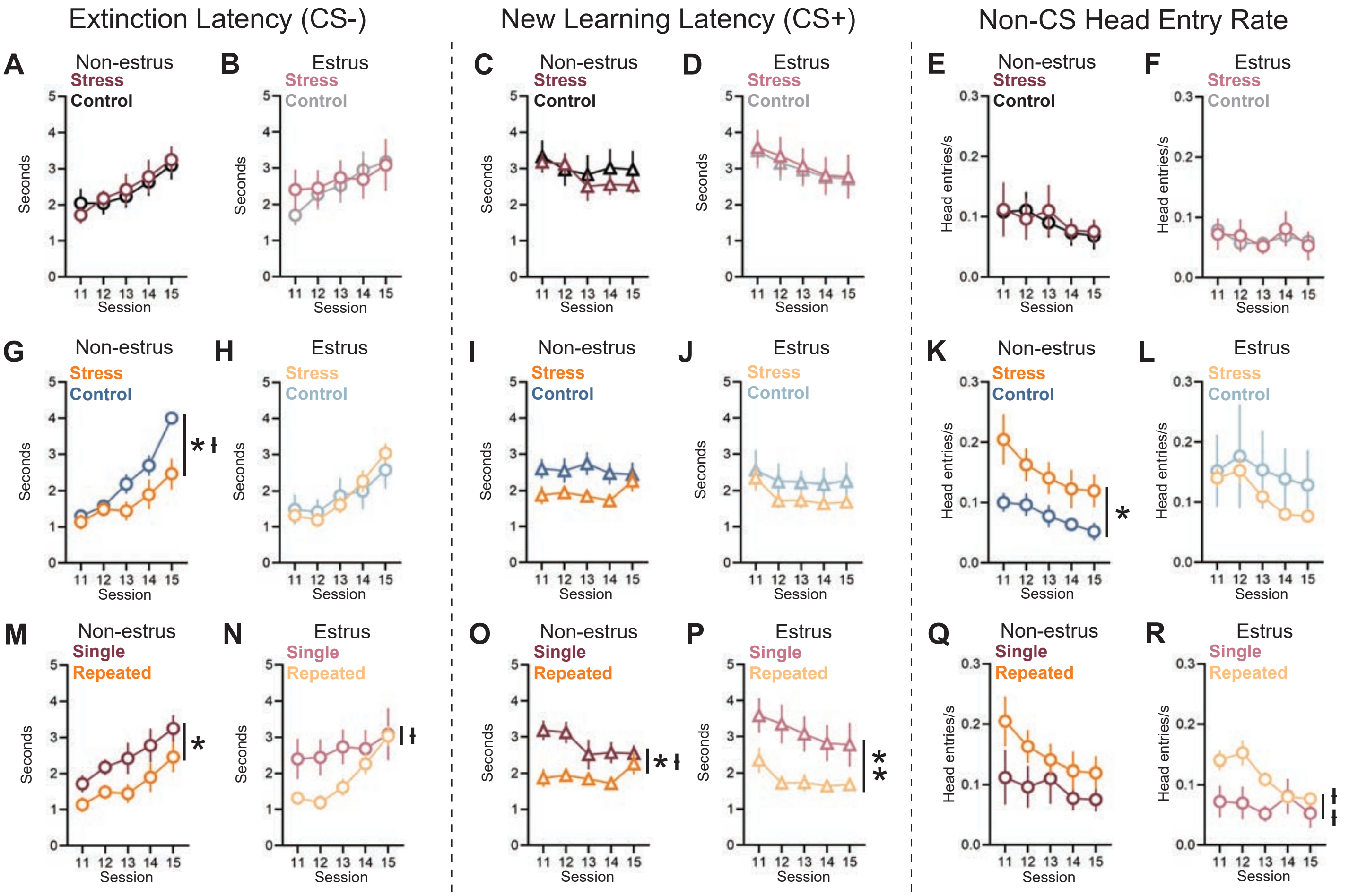


**Supplementary Figure 10:** Latency of rats to food port after onset of cue during contingency change sessions. (A-B) Latency of non-estrus (A) and estrus (B) rats to food port after onset of white noise (extinction) cue does not significantly differ between stress and control. (C-D) Latency of non-estrus (C) and estrus (D) rats to food port after onset of tone (new learning) does not significantly differ between stress and control. (E-F) Non-CS head entry rate did not differ between single stress and single control treatment in both non-estrus (E) and estrus (F) rats. (G) Latency of non-estrus repeated stress rats is faster than that of non-estrus repeated control rats after onset of extinction cue (main effect of stress: $F_{(1,14)}$ = 5.832, * p = 0.0300; interaction of session x stress: $F_{(4,49)}$ = 3.306, ł p = 0.0178). (H) Latency of estrus rats after onset of extinction cue does not significantly differ between repeated stress and repeated control groups. (I-J) Latency of non-estrus (I) and estrus (J) rats after onset of new learning cue does not significantly differ between repeated stress and repeated control groups. (K) Repeated stress non-estrus rats had a greater non-CS head entry rate relative to repeated control non-estrus rats (main effect of stress: $F_{(1,14)}$ = 5.481, * p = 0.0345). (L) Non-CS head entry rates of repeated stress estrus and repeated control estrus rats did not differ. (M) Latency of non-estrus repeated stress rats is faster than that of non-estrus single stress rats after onset of extinction cue (main effect of repeated stress: $F_{(1,10)}$ = 5.660, * p = 0.0387). (N) Latency of estrus repeated stress rats is faster than that of estrus single stress rats after onset of extinction cue (interaction of session x repeated stress: $F_{(4,50)}$ = 2.669, ł p = 0.0428). (O) Latency of non-estrus repeated stress rats is faster than that of non-estrus single stress rats after onset of new learning cue (main effect of repeated stress: $F_{(1,10)}$ = 9.633, * p = 0.0112; interaction of session x repeated stress: $F_{(4,37)}$ = 2.917, ł p = 0.0341). (P) Latency of estrus repeated stress rats is faster than that of estrus single stress rats after onset of new learning cue (main effect of repeated stress: $F_{(1,13)}$ = 9.236, ** p = 0.0095). (Q) Non-CS head entry rate of single and repeated stress non-estrus rats did not differ. (R) Repeated stress estrus rats had greater non-CS head entries relative to single stress estrus rats (interaction of session x repeated stress: $F_{(4,50)}$ = 4.701, łł p = 0.0027).

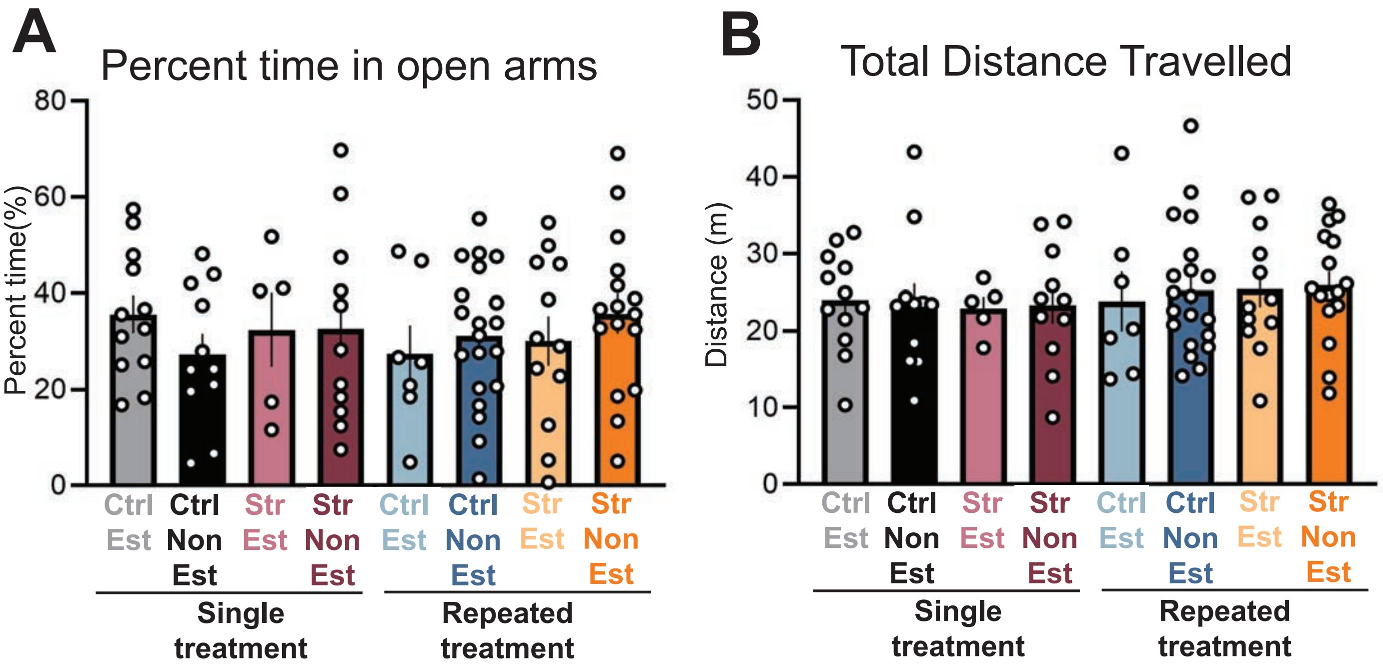


**Supplementary Figure 11:** Estrous cycle and stress treatments did not impact behavior on the zero-maze test. (A) No difference in the percentage of time spent in the open arms between groups. (B) No difference in the total distance rats travelled between groups.

| Table 1 | | | |
|---|---|---|---|
| Figure 1 | | | |
| all single stress females n=34; all single control females n=37; single stress non-estrus n=23; single stress estrus n=11; single control non-estrus n=21; single control estrus n=16 | | | |
| Panel A - experimental timeline | | | |
| Panel B - schematic of trial structure | | | |
| Panel C - Conditioned responding of female rats, not separated by the estrous cycle | | | |
| Two-way mixed effects model | Session | Stress | Two-way interaction |
| | **F (1.522, 99.28) = 85.49, p<0.0001** | F (1, 69) = 0.2933, p=0.5899 | F (9, 587) = 0.3152, p=0.9701 |
| Panel D - Conditioned responding of non-estrus rats | | | |
| Two-way mixed effects model | Session | Stress | Two-way interaction |
| | **F (1.837, 71.66) = 62.16, p<0.0001** | **F (1, 42) = 10.69, p=0.0022** | **F (9, 351) = 1.975, p=0.0412** |
| Post-hoc Tukey's multiple comparisons test | Session | 95.00% CI of diff. | Adjusted P Value |
| | **1** | **-0.2345 to -0.02217** | **0.0191** |
| | **2** | **-0.2625 to -0.003814** | **0.044** |
| | **3** | **-0.3746 to -0.03623** | **0.0188** |
| | 4 | -0.4392 to 0.04203 | 0.1028 |
| | **5** | **-0.5936 to -0.05009** | **0.0223** |
| | **6** | **-0.5245 to -0.08442** | **0.0082** |
| | **7** | **-0.5869 to -0.03000** | **0.0311** |
| | **8** | **-0.7203 to -0.1251** | **0.0069** |
| | **9** | **-0.6575 to -0.1098** | **0.0075** |
| | **10** | **-0.6434 to -0.04099** | **0.0272** |
| Panel E - Conditioned responding of estrus rats | | | |
| Two-way mixed effects model | Session | Stress | Two-way interaction |
| | **F (1.372, 33.24) = 25.77, p<0.0001** | **F (1, 25) = 5.862, p=0.0231** | **F (9, 218) = 2.021, p=0.0382** |
| Post-hoc Tukey's multiple comparisons test | Session | 95.00% CI of diff. | Adjusted P Value |
| | 1 | -0.02687 to 0.1902 | 0.1331 |
| | 2 | -0.003414 to 0.3932 | 0.0538 |
| | 3 | -0.04943 to 0.3973 | 0.1213 |
| | 4 | -0.03441 to 0.6145 | 0.0772 |
| | **5** | **0.06475 to 0.8167** | **0.0241** |
| | **6** | **0.04280 to 0.8303** | **0.0315** |
| | **7** | **0.1503 to 0.8021** | **0.006** |
| | **8** | **0.1457 to 0.8340** | **0.0073** |
| | **9** | **0.06080 to 0.7674** | **0.0236** |
| | **10** | **0.02844 to 0.7880** | **0.0364** |
| Panel F - Latency of female rats, not separated by the estrous cycle | | | |
| Two-way mixed effects model | Session | Stress | Two-way interaction |
| | **F (1.755, 114.5) = 126.5, p<0.0001** | F (1, 69) = 0.02656, p=0.8710 | F (9, 587) = 1.522, p=0.1366 |
| Panel G - Latency of non-estrus rats | | | |
| Two-way mixed effects model | Session | Stress | Two-way interaction |
| | **F (1.664, 64.91) = 76.10, p<0.0001** | F (1, 42) = 0.4319, p=0.5146 | **F (9, 351) = 3.488, p=0.0004** |
| Post-hoc Tukey's multiple comparisons test | Session | 95.00% CI of diff. | Adjusted P Value |
| | **1** | **0.1084 to 1.362** | **0.0226** |
| | 2 | -0.1849 to 0.8962 | 0.1913 |
| | 3 | -0.4291 to 0.5203 | 0.8472 |
| | 4 | -0.5461 to 0.4160 | 0.7863 |
| | 5 | -0.4904 to 0.3057 | 0.6414 |
| | 6 | -0.4043 to 0.4266 | 0.957 |
| | 7 | -0.4852 to 0.3287 | 0.6992 |
| | 8 | -0.3549 to 0.5850 | 0.6216 |
| | 9 | -0.1816 to 0.5699 | 0.3019 |
| | 10 | -0.3927 to 0.4843 | 0.8332 |
| Panel H - Latency of estrus rats | | | |
| Two-way mixed effects model | Session | Stress | Two-way interaction |
| | **F (1.846, 44.71) = 55.65, p<0.0001** | F (1, 25) = 0.2524, p=0.6198 | F (9, 218) = 0.5037, p=0.8710 |
| Panel I - Non-CS head entry rate of female rats, not separated by the estrous cycle | | | |
| Two-way mixed effects model | Session | Stress | Two-way interaction |
| | **F (1.598, 103.9) = 4.244, p=0.0243** | F (1, 69) = 0.0004076, p=0.9840 | F (9, 585) = 1.401, p=0.1840 |
| Panel J - Non-CS head entry rate of non-estrus rats | | | |
| Two-way mixed effects model | Session | Stress | Two-way interaction |
| | F (1.711, 66.72) = 2.787, p=0.0768 | F (1, 42) = 0.3458, p=0.5596 | **F (9, 351) = 2.120, p=0.0273** |
| Post-hoc Tukey's multiple comparisons test | Session | 95.00% CI of diff. | Adjusted P Value |
| | 1 | -0.08510 to 0.01667 | 0.1817 |
| | 2 | -0.04787 to 0.05084 | 0.9517 |
| | 3 | -0.02405 to 0.04051 | 0.6093 |
| | 4 | -0.04674 to 0.04814 | 0.9763 |
| | 5 | -0.01446 to 0.07383 | 0.1815 |
| | 6 | -0.02635 to 0.1028 | 0.2384 |
| | 7 | -0.04841 to 0.08370 | 0.5909 |
| | 8 | -0.05241 to 0.07759 | 0.697 |
| | 9 | -0.06803 to 0.04716 | 0.7144 |
| | 10 | -0.01487 to 0.09601 | 0.1463 |
| Panel K - Non-CS head entry rate of estrus rats | | | |
| Two-way mixed effects model | Session | Stress | Two-way interaction |
| | F (1.351, 32.41) = 1.825, p=0.1852 | F (1, 25) = 0.4582, p=0.5047 | F (9, 216) = 0.8469, p=0.5737 |

| Figure 2 | | | |
|---|---|---|---|
| all repeated stress females n=29; all repeated control females n=27; repeated stress non-estrus n=17; repeated stress estrus n=12; repeated control non-estrus n=20; repeated control estrus n=7 | | | |
| Panel A - experimental timeline | | | |
| Panel B - Conditioned responding of female rats, not separated by the estrous cycle | | | |
| Two-way mixed effects model | Session | Stress | Two-way interaction |
| | **F (2.224, 114.7) = 62.04, p<0.0001** | F (1, 54) = 2.238, p=0.1405 | **F (9, 464) = 2.695, p=0.0046** |
| Post-hoc Tukey's multiple comparisons test | Session | 95.00% CI of diff. | Adjusted P Value |
| | 1 | -0.06389 to 0.1197 | 0.5432 |
| | 2 | -0.1121 to 0.1284 | 0.8922 |
| | 3 | -0.2319 to 0.1199 | 0.5252 |
| | **4** | **-0.5286 to -0.08709** | **0.0073** |
| | **5** | **-0.5659 to -0.08579** | **0.0087** |
| | **6** | **-0.5525 to -0.03224** | **0.0284** |
| | 7 | -0.4623 to 0.1025 | 0.2068 |
| | 8 | -0.2250 to 0.4032 | 0.571 |
| | 9 | -0.4523 to 0.1935 | 0.4242 |
| | 10 | -0.4714 to 0.1974 | 0.4137 |
| Panel C - Conditioned responding of non-estrus rats | | | |
| Two-way mixed effects model | Session | Stress | Two-way interaction |
| | **F (2.545, 85.39) = 41.36, p<0.0001** | F (1, 35) = 3.596, p=0.0662 | **F (9, 302) = 2.461, p=0.0102** |
| Post-hoc Tukey's multiple comparisons test | Session | 95.00% CI of diff. | Adjusted P Value |
| | 1 | -0.08650 to 0.1637 | 0.5347 |
| | 2 | -0.07068 to 0.2308 | 0.2881 |
| | 3 | -0.3425 to 0.1679 | 0.4872 |
| | **4** | **-0.6198 to -0.08251** | **0.0127** |
| | **5** | **-0.7108 to -0.1060** | **0.0096** |
| | **6** | **-0.7620 to -0.1335** | **0.0066** |
| | 7 | -0.6419 to 0.09657 | 0.1426 |
| | 8 | -0.5044 to 0.2796 | 0.5628 |
| | 9 | -0.6902 to 0.1207 | 0.1621 |
| | 10 | -0.7109 to 0.1223 | 0.1593 |
| Panel D - Conditioned responding of estrus rats | | | |
| Two-way mixed effects model | Session | Stress | Two-way interaction |
| | **F (1.641, 26.26) = 23.22, p<0.0001** | F (1, 17) = 0.07020, p=0.7942 | F (9, 144) = 1.230, p=0.2809 |
| Panel E - Latency of female rats, not separated by the estrous cycle | | | |
| Two-way mixed effects model | Session | Stress | Two-way interaction |
| | **F (2.972, 153.2) = 104.0, p<0.0001** | F (1, 54) = 3.638, p=0.0618 | F (9, 464) = 1.208, p=0.2875 |
| Panel F - Latency of non-estrus rats | | | |
| Two-way mixed effects model | Session | Stress | Two-way interaction |
| | **F (3.106, 104.2) = 55.25, p<0.0001** | F (1, 35) = 0.7591, p=0.3896 | F (9, 302) = 0.5651, p=0.8253 |
| Panel G - Latency of estrus rats | | | |
| Two-way mixed effects model | Session | Stress | Two-way interaction |
| | **F (2.303, 36.85) = 53.19, p<0.0001** | **F (1, 17) = 4.725, p=0.0441** | F (9, 144) = 1.387, p=0.1993 |
| Post-hoc Tukey's multiple comparisons test | Session | 95.00% CI of diff. | Adjusted P Value |
| | 1 | -0.5960 to 1.164 | 0.4712 |
| | 2 | -0.06166 to 1.401 | 0.0668 |
| | 3 | -0.3786 to 0.6762 | 0.5304 |
| | 4 | -0.008100 to 1.454 | 0.0519 |
| | 5 | -0.3582 to 1.382 | 0.2042 |
| | 6 | -0.3476 to 1.303 | 0.2095 |
| | 7 | -0.2211 to 1.106 | 0.1563 |
| | 8 | -0.4376 to 0.4994 | 0.8804 |
| | 9 | -0.4728 to 0.7553 | 0.5897 |
| | 10 | -0.4501 to 0.7445 | 0.5649 |
| Panel H - Non-CS head entry rate of female rats, not separated by the estrous cycle | | | |
| Two-way mixed effects model | Session | Stress | Two-way interaction |
| | **F (1.611, 83.08) = 4.512, p=0.0200** | F (1, 54) = 3.410, p=0.0703 | F (9, 464) = 0.5784, p=0.8151 |
| Panel I - Non-CS head entry rate of non-estrus rats | | | |
| Two-way mixed effects model | Session | Stress | Two-way interaction |
| | F (1.886, 63.29) = 2.881, p=0.0665 | **F (1, 35) = 4.285, p=0.0459** | F (9, 302) = 1.080, p=0.3774 |
| Post-hoc Tukey's multiple comparisons test | Session | 95.00% CI of diff. | Adjusted P Value |
| | 1 | -0.09124 to 0.01378 | 0.1425 |
| | **2** | **-0.1135 to -0.01530** | **0.0121** |
| | 3 | -0.05861 to 0.03168 | 0.5471 |
| | 4 | -0.1031 to 0.001071 | 0.0546 |
| | 5 | -0.1174 to 0.007028 | 0.0801 |
| | 6 | -0.08588 to 0.04032 | 0.4677 |
| | 7 | -0.1012 to 0.03070 | 0.2838 |
| | 8 | -0.1291 to 0.02272 | 0.1609 |
| | 9 | -0.1290 to 0.01337 | 0.1067 |
| | 10 | -0.1315 to 0.008194 | 0.0811 |
| Panel J - Non-CS head entry rate of estrus rats | | | |
| Two-way mixed effects model | Session | Stress | Two-way interaction |
| | F (1.275, 20.40) = 1.969, p=0.1744 | F (1, 17) = 0.4238, p=0.5238 | F (9, 144) = 0.6480, p=0.7544 |

| Figure 3 | | | |
|---|---|---|---|
| single stress non-estrus n=23; repeated stress non-estrus n=17; single stress estrus n=11; repeated stress estrus n=12 | | | |
| Panel A - experimental timeline | | | |
| Panel B - Conditioned responding of non-estrus rats | | | |
| Two-way mixed effects model | Session | Repeated treatment | Two-way interaction |
| | **F (1.869, 63.97) = 64.85, p<0.0001** | **F (1, 38) = 5.312, p=0.0267** | **F (9, 308) = 2.198, p=0.0221** |
| Post-hoc Tukey's multiple comparisons test | Session | 95.00% CI of diff. | Adjusted P Value |
| | 1 | -0.1068 to 0.1239 | 0.8813 |
| | 2 | -0.2871 to 0.03215 | 0.1142 |
| | 3 | -0.5206 to 0.01760 | 0.0659 |
| | **4** | **-0.7626 to -0.1297** | **0.0071** |
| | **5** | **-0.6938 to -0.01536** | **0.041** |
| | **6** | **-0.7175 to -0.1916** | **0.0013** |
| | 7 | -0.6731 to 0.02431 | 0.0672 |
| | 8 | -0.6024 to 0.1645 | 0.2524 |
| | 9 | -0.6576 to 0.08035 | 0.1202 |
| | 10 | -0.6315 to 0.1180 | 0.1708 |
| Panel C - Conditioned responding of estrus rats | | | |
| Two-way mixed effects model | Session | Repeated treatment | Two-way interaction |
| | **F (1.510, 29.54) = 24.84, p<0.0001** | **F (1, 21) = 12.41, p=0.0020** | F (9, 176) = 1.909, p=0.0534 |
| Post-hoc Tukey's multiple comparisons test | Session | 95.00% CI of diff. | Adjusted P Value |
| | **1** | **-0.1812 to -0.03405** | **0.0062** |
| | **2** | **-0.4683 to -0.1602** | **0.0004** |
| | **3** | **-0.5794 to -0.1210** | **0.0047** |
| | **4** | **-0.7955 to -0.1117** | **0.0124** |
| | **5** | **-0.8900 to -0.2300** | **0.0027** |
| | **6** | **-0.8708 to -0.2541** | **0.0013** |
| | **7** | **-0.9058 to -0.2012** | **0.0041** |
| | 8 | -0.6678 to 0.07821 | 0.1121 |
| | 9 | -0.8242 to 0.02824 | 0.0653 |
| | 10 | -0.9275 to 0.09144 | 0.1007 |
| Panel D - Latency of non-estrus rats | | | |
| Two-way mixed effects model | Session | Repeated treatment | Two-way interaction |
| | **F (1.854, 63.44) = 51.94, p<0.0001** | **F (1, 38) = 8.370, p=0.0063** | F (9, 308) = 1.006, p=0.4348 |
| Post-hoc Tukey's multiple comparisons test | Session | 95.00% CI of diff. | Adjusted P Value |
| | 1 | -0.3568 to 0.9182 | 0.3783 |
| | **2** | **0.1806 to 1.211** | **0.0095** |
| | **3** | **0.1046 to 0.9676** | **0.0163** |
| | **4** | **0.2637 to 1.095** | **0.0021** |
| | **5** | **0.1549 to 0.9253** | **0.0074** |
| | **6** | **0.3376 to 0.9599** | **0.0002** |
| | **7** | **0.06352 to 0.8448** | **0.024** |
| | 8 | -0.03586 to 0.7320 | 0.0739 |
| | **9** | **0.1142 to 0.7631** | **0.0097** |
| | **10** | **0.1237 to 0.8468** | **0.0103** |
| Panel E - Latency of estrus rats | | | |
| Two-way mixed effects model | Session | Repeated treatment | Two-way interaction |
| | **F (2.015, 39.41) = 75.68, p<0.0001** | **F (1, 21) = 14.63, p=0.0010** | F (9, 176) = 1.188, p=0.3054 |
| Post-hoc Tukey's multiple comparisons test | Session | 95.00% CI of diff. | Adjusted P Value |
| | 1 | -0.02835 to 1.243 | 0.0596 |
| | **2** | **0.3178 to 1.462** | **0.0056** |
| | 3 | -0.09029 to 0.9218 | 0.0989 |
| | **4** | **0.1477 to 0.9071** | **0.0105** |
| | **5** | **0.1816 to 0.7767** | **0.0035** |
| | **6** | **0.05855 to 0.6859** | **0.0236** |
| | **7** | **0.1376 to 0.7652** | **0.0087** |
| | **8** | **0.1886 to 0.7845** | **0.0032** |
| | **9** | **0.1486 to 0.7996** | **0.0079** |
| | **10** | **0.1412 to 0.9731** | **0.0137** |
| Panel F - Non-CS head entry rate of non-estrus rats | | | |
| Two-way mixed effects model | Session | Repeated treatment | Two-way interaction |
| | **F (1.806, 61.80) = 3.846, p=0.0305** | **F (1, 38) = 16.45, p=0.0002** | F (9, 308) = 1.234, p=0.2733 |
| Post-hoc Tukey's multiple comparisons test | Session | 95.00% CI of diff. | Adjusted P Value |
| | **1** | **-0.1241 to -0.006620** | **0.0302** |
| | **2** | **-0.1501 to -0.04849** | **0.0004** |
| | **3** | **-0.1034 to -0.01465** | **0.0108** |
| | **4** | **-0.1411 to -0.02887** | **0.0041** |
| | **5** | **-0.1777 to -0.05737** | **0.0004** |
| | **6** | **-0.1689 to -0.04337** | **0.0016** |
| | **7** | **-0.1522 to -0.004962** | **0.0372** |
| | **8** | **-0.1697 to -0.01185** | **0.0258** |
| | **9** | **-0.1714 to -0.01921** | **0.016** |
| | **10** | **-0.1855 to -0.04673** | **0.002** |
| Panel G - Non-CS head entry rate of estrus rats | | | |
| Two-way mixed effects model | Session | Repeated treatment | Two-way interaction |
| | F (1.250, 24.44) = 2.611, p=0.1126 | F (1, 21) = 0.6131, p=0.4424 | F (9, 176) = 0.6573, p=0.7466 |

| Figure 4 | | | |
|---|---|---|---|
| single control non-estrus n=21; single control estrus n=16; single stress non-estrus n=23; single stress estrus n=11; repeated control non-estrus n=20; repeated control estrus n=7; repeated stress non-estrus n=17; repeated stress estrus n=12 | | | |
| Panel A - Conditioned responding of single control rats | | | |
| Two-way mixed effects model | Session | Stage | Two-way interaction |
| | **F (1.550, 53.92) = 45.91, p<0.0001** | **F (1, 35) = 6.932, p=0.0125** | F (9, 313) = 1.253, p=0.2619 |
| Post-hoc Tukey's multiple comparisons test | Session | 95.00% CI of diff. | Adjusted P Value |
| | 1 | -0.02065 to 0.2065 | 0.1048 |
| | **2** | **0.04102 to 0.3956** | **0.0185** |
| | **3** | **0.01169 to 0.4122** | **0.039** |
| | 4 | -0.009748 to 0.6149 | 0.0569 |
| | **5** | **0.03047 to 0.7735** | **0.0356** |
| | 6 | -0.01637 to 0.7543 | 0.0595 |
| | 7 | 0.05023 to 0.6750 | 0.0248 |
| | 8 | -0.0006522 to 0.6768 | 0.0504 |
| | 9 | -0.06844 to 0.5372 | 0.1237 |
| | 10 | -0.07690 to 0.5977 | 0.1252 |
| Panel B - Conditioned responding of single stress rats | | | |
| Two-way mixed effects model | Session | Stage | Two-way interaction |
| | **F (1.651, 46.96) = 34.42, p<0.0001** | **F (1, 32) = 9.240, p=0.0047** | **F (9, 256) = 3.959, p<0.0001** |
| Post-hoc Tukey's multiple comparisons test | Session | 95.00% CI of diff. | Adjusted P Value |
| | **1** | **-0.2174 to -0.01658** | **0.0238** |
| | 2 | -0.2698 to 0.05021 | 0.1711 |
| | 3 | -0.3653 to 0.03051 | 0.0944 |
| | 4 | -0.4442 to 0.07206 | 0.1518 |
| | **5** | **-0.6392 to -0.08199** | **0.0133** |
| | **6** | **-0.6101 to -0.1339** | **0.0034** |
| | **7** | **-0.7165 to -0.1276** | **0.0066** |
| | **8** | **-0.8790 to -0.2700** | **0.0006** |
| | **9** | **-0.8944 to -0.2324** | **0.0019** |
| | **10** | **-0.8409 to -0.1391** | **0.0087** |
| Panel C - Conditioned responding of repeated control rats | | | |
| Two-way mixed effects model | Session | Stage | Two-way interaction |
| | **F (2.498, 60.50) = 18.46, p<0.0001** | F (1, 25) = 0.02672, p=0.8715 | F (9, 218) = 0.4920, p=0.8791 |
| Panel D - Conditioned responding of repeated stress rats | | | |
| Two-way mixed effects model | Session | Stage | Two-way interaction |
| | **F (1.869, 47.34) = 40.37, p<0.0001** | F (1, 27) = 2.028, p=0.1658 | F (9, 228) = 1.734, p=0.0824 |

| Figure 5 | | | |
|---|---|---|---|
| single stress non-estrus n=6; single stress estrus n=5; single control non-estrus n=8; single control estrus n=6; repeated stress non-estrus n=6; repeated stress estrus n=10; repeated control non-estrus n=8; repeated control estrus n=8 | | | |
| Panel A - Experimental timeline | | | |
| Panel B - Extinction of single treatment non-estrus rats | | | |
| Two-way mixed effects model | Session | Stress | Two-way interaction |
| | **F (1.661, 19.10) = 10.01, p=0.0017** | F (1, 12) = 1.664, p=0.2213 | F (4, 46) = 1.059, p=0.3874 |
| Panel C - Extinction of single treatment estrus rats | | | |
| Two-way mixed effects model | Session | Stress | Two-way interaction |
| | **F (2.168, 18.97) = 8.375, p=0.0021** | F (1, 9) = 0.1007, p=0.7583 | F (4, 35) = 0.4722, p=0.7558 |
| Panel D - New learning of single treatment non-estrus rats | | | |
| Two-way mixed effects model | Session | Stress | Two-way interaction |
| | **F (2.567, 29.52) = 9.016, p=0.0004** | F (1, 12) = 3.656, p=0.0800 | F (4, 46) = 1.920, p=0.1232 |
| Panel E - New learning of single treatment estrus rats | | | |
| Two-way mixed effects model | Session | Stress | Two-way interaction |
| | F (1.576, 13.79) = 3.939, p=0.0524 | F (1, 9) = 2.169, p=0.1749 | F (4, 35) = 0.3309, p=0.8553 |
| Panel F - Experimental timeline | | | |
| Panel G - Extinction of repeated treatment non-estrus rats | | | |
| Two-way mixed effects model | Session | Stress | Two-way interaction |
| | **F (1.462, 17.91) = 16.48, p=0.0002** | F (1, 14) = 0.003322, p=0.9549 | F (4, 49) = 0.6946, p=0.5993 |
| Panel H - Extinction of repeated treatment estrus rats | | | |
| Two-way mixed effects model | Session | Stress | Two-way interaction |
| | **F (3.046, 44.17) = 26.00, p<0.0001** | F (1, 15) = 0.4512, p=0.5120 | F (4, 58) = 0.2308, p=0.9200 |
| Panel I - New learning of repeated treatment non-estrus rats | | | |
| Two-way mixed effects model | Session | Stress | Two-way interaction |
| | **F (2.030, 24.87) = 4.478, p=0.0214** | **F (1, 14) = 5.683, p=0.0318** | F (4, 49) = 0.8540, p=0.4981 |
| Post-hoc Tukey's multiple comparisons test | Session | 95.00% CI of diff. | Adjusted P Value |
| | 11 | -0.7534 to 0.1145 | 0.124 |
| | 12 | -0.6291 to 0.06892 | 0.1013 |
| | **13** | **-0.7748 to -0.05639** | **0.0289** |
| | 14 | -0.9043 to 0.07856 | 0.0891 |
| | 15 | -0.6938 to 0.6402 | 0.9227 |
| Panel J - New learning of repeated treatment estrus rats | | | |
| Two-way mixed effects model | Session | Stress | Two-way interaction |
| | **F (1.969, 28.55) = 8.206, p=0.0016** | F (1, 15) = 2.133, p=0.1648 | **F (4, 58) = 2.715, p=0.0384** |
| Post-hoc Tukey's multiple comparisons test | Session | 95.00% CI of diff. | Adjusted P Value |
| | 11 | -0.3451 to 0.4157 | 0.8443 |
| | 12 | -0.5625 to 0.2665 | 0.4534 |
| | 13 | -0.6698 to 0.006443 | 0.0539 |
| | **14** | **-0.7362 to -0.0005295** | **0.0497** |
| | 15 | -0.9609 to 0.1787 | 0.1477 |
| Panel K - Experimental timeline | | | |
| Panel L - Extinction of stressed non-estrus rats | | | |
| Two-way mixed effects model | Session | Repeated treatment | Two-way interaction |
| | **F (1.420, 13.14) = 9.287, p=0.0055** | F (1, 10) = 0.002503, p=0.9611 | F (4, 37) = 0.7744, p=0.5489 |
| Panel M - Extinction of stressed estrus rats | | | |
| Two-way mixed effects model | Session | Repeated treatment | Two-way interaction |
| | **F (2.091, 26.14) = 9.240, p=0.0008** | F (1, 13) = 0.004139, p=0.9497 | F (4, 50) = 2.557, p=0.0500 |
| Panel N - New learning of stressed non-estrus rats | | | |
| Two-way mixed effects model | Session | Repeated treatment | Two-way interaction |
| | **F (1.773, 16.40) = 4.898, p=0.0246** | **F (1, 10) = 5.644, p=0.0389** | F (4, 37) = 0.3487, p=0.8432 |
| Post-hoc Tukey's multiple comparisons test | Session | 95.00% CI of diff. | Adjusted P Value |
| | 11 | -0.8428 to 0.05082 | 0.0751 |
| | **12** | **-0.7180 to -0.02868** | **0.0376** |
| | 13 | -0.7790 to 0.01742 | 0.0587 |
| | 14 | -0.8544 to 0.1130 | 0.1128 |
| | 15 | -0.8096 to 0.5122 | 0.5763 |
| Panel O - New learning of stressed estrus rats | | | |
| Two-way mixed effects model | Session | Repeated treatment | Two-way interaction |
| | **F (2.125, 26.56) = 9.675, p=0.0006** | **F (1, 13) = 11.87, p=0.0043** | **F (4, 50) = 3.133, p=0.0224** |
| Post-hoc Tukey's multiple comparisons test | Session | 95.00% CI of diff. | Adjusted P Value |
| | 11 | -0.5170 to 0.03216 | 0.0785 |
| | **12** | **-0.8271 to -0.2465** | **0.0016** |
| | **13** | **-0.9339 to -0.2565** | **0.0023** |
| | **14** | **-0.9751 to -0.3577** | **0.0005** |
| | **15** | **-0.9777 to -0.4192** | **0.0003** |

| Supplementary Figure 1 | | | |
|---|---|---|---|
| single stress proestrus n=8; single control proestrus n=8; single stress estrus n=11; single control estrus n=16; single stress metestrus n=6; single control metestrus n=6; single stress diestrus n=9; single control diestrus n=7 | | | |
| Panel A - Conditioned responding of estrus rats | | | |
| Two-way mixed effects model | Session | Stress | Two-way interaction |
| | **F (1.372, 33.24) = 25.77, p<0.0001** | **F (1, 25) = 5.862, p=0.0231** | **F (9, 218) = 2.021, p=0.0382** |
| Post-hoc Tukey's multiple comparisons test | Session | 95.00% CI of diff. | Adjusted P Value |
| | 1 | -0.02687 to 0.1902 | 0.1331 |
| | 2 | -0.003414 to 0.3932 | 0.0538 |
| | 3 | -0.04943 to 0.3973 | 0.1213 |
| | 4 | -0.03441 to 0.6145 | 0.0772 |
| | **5** | **0.06475 to 0.8167** | **0.0241** |
| | **6** | **0.04280 to 0.8303** | **0.0315** |
| | **7** | **0.1503 to 0.8021** | **0.006** |
| | **8** | **0.1457 to 0.8340** | **0.0073** |
| | **9** | **0.06080 to 0.7674** | **0.0236** |
| | **10** | **0.02844 to 0.7880** | **0.0364** |
| Panel B - Conditioned responding of metestrus rats | | | |
| Two-way mixed effects model | Session | Stress | Two-way interaction |
| | **F (3.750, 30.83) = 15.12, p<0.0001** | **F (1, 10) = 7.311, p=0.0222** | F (9, 74) = 0.9293, p=0.5051 |
| Post-hoc Tukey's multiple comparisons test | Session | 95.00% CI of diff. | Adjusted P Value |
| | 1 | -0.3759 to 0.1826 | 0.4576 |
| | **2** | **-0.4283 to -0.01037** | **0.0414** |
| | 3 | -0.5993 to 0.1113 | 0.1555 |
| | 4 | -0.5386 to 0.3920 | 0.7228 |
| | 5 | -0.8406 to 0.1399 | 0.1143 |
| | 6 | -0.9832 to 0.2932 | 0.1978 |
| | 7 | -0.8682 to 0.5268 | 0.5085 |
| | 8 | -0.7348 to 0.2735 | 0.3122 |
| | **9** | **-1.133 to -0.1186** | **0.0224** |
| | 10 | -1.504 to 0.3049 | 0.1526 |
| Panel C - Conditioned responding of diestrus rats | | | |
| Two-way mixed effects model | Session | Stress | Two-way interaction |
| | **F (1.534, 19.77) = 22.98, p<0.0001** | **F (1, 14) = 7.312, p=0.0171** | F (9, 116) = 1.864, p=0.0642 |
| Post-hoc Tukey's multiple comparisons test | Session | 95.00% CI of diff. | Adjusted P Value |
| | **1** | **-0.4311 to -0.04862** | **0.0184** |
| | 2 | -0.4573 to 0.07434 | 0.14 |
| | **3** | **-0.6510 to -0.02541** | **0.0368** |
| | 4 | -0.9373 to 0.02376 | 0.0601 |
| | **5** | **-1.154 to -0.006617** | **0.048** |
| | **6** | **-0.7557 to -0.04690** | **0.0301** |
| | **7** | **-1.048 to -0.03385** | **0.0387** |
| | **8** | **-1.396 to -0.1827** | **0.016** |
| | **9** | **-1.040 to -0.08668** | **0.0249** |
| | 10 | -0.6397 to 0.2623 | 0.3768 |
| Panel D - Conditioned responding of proestrus rats | | | |
| Two-way mixed effects model | Session | Stress | Two-way interaction |
| | **F (1.586, 22.03) = 30.98, p<0.0001** | F (1, 14) = 1.525, p=0.2371 | **F (9, 125) = 1.957, p=0.0498** |
| Post-hoc Tukey's multiple comparisons test | Session | 95.00% CI of diff. | Adjusted P Value |
| | 1 | -0.1300 to 0.05798 | 0.4178 |
| | 2 | -0.2373 to 0.2078 | 0.8854 |
| | 3 | -0.2741 to 0.1921 | 0.7096 |
| | 4 | -0.3141 to 0.2421 | 0.7852 |
| | 5 | -0.4298 to 0.2398 | 0.5456 |
| | 6 | -0.5636 to 0.06464 | 0.1104 |
| | 7 | -0.6311 to 0.1641 | 0.2263 |
| | 8 | -0.5993 to 0.1633 | 0.2403 |
| | 9 | -0.5846 to 0.2206 | 0.3479 |
| | **10** | **-0.9016 to -0.004803** | **0.048** |

| Supplementary Figure 2 | | | |
|---|---|---|---|
| Panel A - Experimental timeline | | | |
| Panel B - Experimental timeline | | | |
| Panel C - Conditioned responding of non-estrus rats | | | |
| Three-way mixed-effects model | Session | Repeated treatment | Stress |
| | **F (2.282, 165.6) = 98.36, p<0.0001** | **F (1, 77) = 18.89, p<0.0001** | **F (1, 77) = 12.68, p=0.0006** |
| | | Session X Repeated treatment | Session X Stress |
| | | F (9, 653) = 1.812, p=0.0629 | **F (9, 653) = 3.466, p=0.0003** |
| | | Repeated treatment X Stress | Three-way interaction |
| | | F (1, 77) = 0.3469, p=0.5576 | **F (9, 653) = 1.399, p=0.1846** |
| Panel D - Conditioned responding of estrus rats | | | |
| Three-way mixed-effects model | Session | Repeated treatment | Stress |
| | **F (1.490, 59.93) = 49.10, p<0.0001** | **F (1, 42) = 4.161, p=0.0477** | F (1, 42) = 1.930, p=0.1721 |
| | | Session X Repeated treatment | Session X Stress |
| | | F (9, 362) = 0.9186, p=0.5088 | **F (9, 362) = 1.981, p=0.0405** |
| | | Repeated treatment X Stress | Three-way interaction |
| | | F (1, 42) = 3.183, p=0.0816 | F (9, 362) = 1.204, p=0.2913 |
| Panel E - Latency of non-estrus rats | | | |
| Three-way mixed-effects model | Session | Repeated treatment | Stress |
| | **F (2.094, 151.9) = 127.4, p<0.0001** | **F (1, 77) = 19.82, p<0.0001** | F (1, 77) = 0.9297, p=0.3379 |
| | | Session X Repeated treatment | Session X Stress |
| | | **F (9, 653) = 2.542, p=0.0071** | F (9, 653) = 1.353, p=0.2059 |
| | | Repeated treatment X Stress | Three-way interaction |
| | | F (1, 77) = 0.01233, p=0.9119 | **F (9, 653) = 2.640, p=0.00522** |
| Panel F - Latency of estrus rats | | | |
| Three-way mixed-effects model | Session | Repeated treatment | Stress |
| | **F (2.018, 81.17) = 98.29, p<0.0001** | F (1, 42) = 2.932, p=0.0942 | F (1, 42) = 1.216, p=0.2765 |
| | | Session X Repeated treatment | Session X Stress |
| | | F (9, 362) = 0.9006, p=0.5248 | F (9, 362) = 0.9458, p=0.4851 |
| | | Repeated treatment X Stress | Three-way interaction |
| | | F (1, 42) = 3.219, p=0.0800 | F (9, 362) = 0.7424, p=0.6699 |
| Panel G - Non-CS head entry rate of non-estrus rats | | | |
| Three-way mixed-effects model | Session | Repeated treatment | Stress |
| | **F (1.830, 132.7) = 5.133, p=0.0088** | **F (1, 77) = 17.13, p<0.0001** | F (1, 77) = 1.304, p=0.2570 |
| | | Session X Repeated treatment | Session X Stress |
| | | F (9, 653) = 0.6199, p=0.7806 | F (9, 653) = 1.534, p=0.1320 |
| | | Repeated treatment X Stress | Three-way interaction |
| | | F (1, 77) = 3.730, p=0.0571 | F (9, 653) = 1.563, p=0.1226 |
| Panel H - Non-CS head entry rate of estrus rats | | | |
| Three-way mixed-effects model | Session | Repeated treatment | Stress |
| | F (1.356, 54.25) = 3.327, p=0.0610 | F (1, 42) = 1.371, p=0.2483 | F (1, 42) = 0.8753, p=0.3548 |
| | | Session X Repeated treatment | Session X Stress |
| | | F (9, 360) = 0.5037, p=0.8717 | F (9, 360) = 0.9635, p=0.4701 |
| | | Repeated treatment X Stress | Three-way interaction |
| | | F (1, 42) = 0.006754, p=0.9349 | F (9, 360) = 0.4913, p=0.8803 |

| Supplementary Figure 3 | | | |
|---|---|---|---|
| single control non-estrus n=21; repeated control non-estrus n=20; single control estrus n=16; repeated control estrus n=7 | | | |
| Panel A - experimental timeline | | | |
| Panel B - Conditioned responding of non-estrus rats | | | |
| Two-way mixed effects model | Session | Repeated treatment | Two-way interaction |
| | **F (2.615, 100.2) = 36.05, p<0.0001** | **F (1, 39) = 17.25, p=0.0002** | F (9, 345) = 1.032, p=0.4140 |
| Post-hoc Tukey's multiple comparisons test | Session | 95.00% CI of diff. | Adjusted P Value |
| | **1** | **-0.2753 to -0.04140** | **0.0095** |
| | **2** | **-0.4584 to -0.2230** | **<0.0001** |
| | **3** | **-0.5122 to -0.2270** | **<0.0001** |
| | **4** | **-0.4581 to -0.1291** | **0.0009** |
| | **5** | **-0.4884 to -0.04768** | **0.019** |
| | **6** | **-0.5926 to -0.02983** | **0.0314** |
| | **7** | **-0.6653 to -0.05511** | **0.0223** |
| | **8** | **-0.8386 to -0.2200** | **0.0015** |
| | **9** | **-0.7125 to -0.06265** | **0.0211** |
| | 10 | -0.6599 to 0.05064 | 0.0902 |
| Panel C - Conditioned responding of estrus rats | | | |
| Two-way mixed effects model | Session | Repeated treatment | Two-way interaction |
| | **F (1.427, 29.50) = 25.79, p<0.0001** | F (1, 21) = 0.02182, p=0.8840 | F (9, 186) = 0.5037, p=0.8707 |
| Panel D - Latency of non-estrus rats | | | |
| Two-way mixed effects model | Session | Repeated treatment | Two-way interaction |
| | **F (2.281, 87.44) = 77.69, p<0.0001** | **F (1, 39) = 11.79, p=0.0014** | **F (9, 345) = 4.314, p<0.0001** |
| Post-hoc Tukey's multiple comparisons test | Session | 95.00% CI of diff. | Adjusted P Value |
| | **1** | **0.4658 to 1.495** | **0.0004** |
| | **2** | **0.6385 to 1.412** | **<0.0001** |
| | **3** | **0.2917 to 1.015** | **0.0009** |
| | **4** | **0.08496 to 0.8243** | **0.0177** |
| | 5 | -0.2593 to 0.6281 | 0.4047 |
| | **6** | **0.005909 to 0.8676** | **0.0471** |
| | 7 | -0.1767 to 0.5766 | 0.2892 |
| | **8** | **0.05426 to 0.9277** | **0.029** |
| | **9** | **0.1614 to 0.8422** | **0.0051** |
| | **10** | **0.04762 to 0.7970** | **0.0286** |
| Panel E - Latency of estrus rats | | | |
| Two-way mixed effects model | Session | Repeated treatment | Two-way interaction |
| | **F (1.863, 38.50) = 34.70, p<0.0001** | F (1, 21) = 0.002417, p=0.9613 | F (9, 186) = 0.5976, p=0.7980 |
| Panel F - Non-CS head entry rate of non-estrus rats | | | |
| Two-way mixed effects model | Session | Repeated treatment | Two-way interaction |
| | F (1.778, 68.17) = 2.565, p=0.0904 | F (1, 39) = 2.741, p=0.1058 | F (9, 345) = 0.8477, p=0.5726 |
| Panel G - Non-CS head entry rate of estrus rats | | | |
| Two-way mixed effects model | Session | Repeated treatment | Two-way interaction |
| | F (1.478, 30.22) = 1.560, p=0.2271 | F (1, 21) = 0.8985, p=0.3540 | F (9, 184) = 0.2873, p=0.9776 |

| | | | |
|---|---|---|---|
| Supplementary Figure 4 | | | |
| repeated control non-estrus n=16; repeated stress non-estrus n=21; repeated control estrus n=11; repeated stress estrus n=8 | | | |
| Panel A - Conditioned responding - non-estrus | | | |
| Two-way mixed effects model | Session | Stress | Two-way interaction |
| | **F (2.202, 73.16) = 40.00, p<0.0001** | F (1, 35) = 1.811, p=0.1870 | **F (9, 299) = 2.793, p=0.0037** |
| Post-hoc Tukey's multiple comparisons test | Session | 95.00% CI of diff. | Adjusted P Value |
| | 1 | -0.08960 to 0.1806 | 0.4913 |
| | 2 | -0.1026 to 0.1941 | 0.5344 |
| | 3 | -0.3413 to 0.1132 | 0.315 |
| | **4** | **-0.5359 to -0.008229** | **0.0437** |
| | **5** | **-0.6943 to -0.05707** | **0.0222** |
| | **6** | **-0.7660 to -0.08412** | **0.0165** |
| | 7 | -0.6604 to 0.04188 | 0.0823 |
| | 8 | -0.2463 to 0.4802 | 0.5162 |
| | 9 | -0.5161 to 0.3041 | 0.6017 |
| | 10 | -0.6100 to 0.2656 | 0.4274 |
| Panel B - Conditioned responding - estrus | | | |
| Two-way mixed effects model | Session | Stress | Two-way interaction |
| | **F (1.973, 32.23) = 21.73, p<0.0001** | F (1, 17) = 0.2695, p=0.6104 | F (9, 147) = 0.8345, p=0.5854 |
| Panel C - Conditioned responding - control | | | |
| Two-way mixed effects model | Session | Stage | Two-way interaction |
| | **F (2.423, 58.70) = 23.49, p<0.0001** | F (1, 25) = 0.05071, p=0.8237 | F (9, 218) = 1.593, p=0.1186 |
| Panel D - Conditioned responding - stress | | | |
| Two-way mixed effects model | Session | Stage | Two-way interaction |
| | **F (1.786, 45.23) = 31.68, p<0.0001** | F (1, 27) = 0.3755, p=0.5451 | F (9, 228) = 0.4785, p=0.8883 |
| Panel E - Latency - non-estrus | | | |
| Two-way mixed effects model | Session | Stress | Two-way interaction |
| | **F (3.152, 104.7) = 58.64, p<0.0001** | F (1, 35) = 0.2333, p=0.6321 | F (9, 299) = 0.7155, p=0.6947 |
| Panel F - Latency - estrus | | | |
| Two-way mixed effects model | Session | Stress | Two-way interaction |
| | **F (2.036, 33.25) = 52.42, p<0.0001** | **F (1, 17) = 4.457, p=0.0499** | F (9, 147) = 0.9090, p=0.5192 |
| Post-hoc Tukey's multiple comparisons test | Session | 95.00% CI of diff. | Adjusted P Value |
| | 1 | -0.3130 to 0.7610 | 0.391 |
| | 2 | -0.03623 to 0.9220 | 0.0674 |
| | 3 | -0.1666 to 0.6337 | 0.2348 |
| | **4** | **0.1203 to 1.120** | **0.0188** |
| | **5** | **0.02351 to 1.162** | **0.0425** |
| | 6 | -0.06590 to 0.9928 | 0.0809 |
| | **7** | **0.01060 to 0.8398** | **0.0452** |
| | 8 | -0.1268 to 0.6229 | 0.1788 |
| | **9** | **0.01979 to 0.8813** | **0.0415** |
| | 10 | -0.03151 to 0.8471 | 0.0665 |
| Panel G - Latency - control | | | |
| Two-way mixed effects model | Session | Stage | Two-way interaction |
| | **F (2.677, 64.84) = 35.72, p<0.0001** | F (1, 25) = 1.792, p=0.1927 | F (9, 218) = 0.8153, p=0.6026 |
| Panel H - Latency - stress | | | |
| Two-way mixed effects model | Session | Stage | Two-way interaction |
| | **F (2.719, 68.89) = 62.09, p<0.0001** | F (1, 27) = 1.856, p=0.1843 | F (9, 228) = 0.4299, p=0.9182 |
| Panel I - Non-CS head entry rate - non-estrus | | | |
| Two-way mixed effects model | Session | Stress | Two-way interaction |
| | **F (1.696, 56.36) = 4.251, p=0.0244** | F (1, 35) = 0.6912, p=0.4114 | F (9, 299) = 0.6941, p=0.7142 |
| Panel J - Non-CS head entry rate - estrus | | | |
| Two-way mixed effects model | Session | Stress | Two-way interaction |
| | F (1.394, 22.76) = 1.336, p=0.2731 | F (1, 17) = 3.427, p=0.0816 | **F (9, 147) = 2.085, p=0.0343** |
| Post-hoc Tukey's multiple comparisons test | Session | 95.00% CI of diff. | Adjusted P Value |
| | 1 | -0.08887 to 0.006562 | 0.0859 |
| | 2 | -0.1413 to 0.01370 | 0.0974 |
| | 3 | -0.08163 to 0.08273 | 0.9886 |
| | 4 | -0.1359 to 0.06119 | 0.4232 |
| | 5 | -0.2283 to 0.02803 | 0.1135 |
| | 6 | -0.2090 to 0.02909 | 0.1254 |
| | 7 | -0.2139 to 0.03228 | 0.134 |
| | 8 | -0.2212 to 0.05571 | 0.2067 |
| | 9 | -0.2426 to 0.04844 | 0.1639 |
| | 10 | -0.1869 to 0.03733 | 0.166 |
| Panel K - Non-CS head entry rate - control | | | |
| Two-way mixed effects model | Session | Stage | Two-way interaction |
| | F (1.566, 37.93) = 2.809, p=0.0845 | F (1, 25) = 0.7984, p=0.3801 | F (9, 218) = 1.429, p=0.1769 |
| Panel L - Non-CS head entry rate - stress | | | |
| Two-way mixed effects model | Session | Stage | Two-way interaction |
| | F (1.556, 39.43) = 2.026, p=0.1543 | F (1, 27) = 0.5295, p=0.4731 | F (9, 228) = 1.715, p=0.0865 |

| Supplementary Figure 6 | | | |
|---|---|---|---|
| single control non-estrus n=21; single control estrus n=16; single stress non-estrus n=23; single stress estrus n=11; repeated control non-estrus n=20; repeated control estrus n=7; repeated stress non-estrus n=17; repeated stress estrus n=12 | | | |
| Panel A - Latency of single control rats | | | |
| Two-way mixed effects model | Session | Stage | Two-way interaction |
| | **F (1.751, 60.89) = 82.95, p<0.0001** | F (1, 35) = 1.911, p=0.1756 | F (9, 313) = 1.312, p=0.2297 |
| Panel B - Latency of single stress rats | | | |
| Two-way mixed effects model | Session | Stage | Two-way interaction |
| | **F (1.639, 46.62) = 45.76, p<0.0001** | F (1, 32) = 0.04403, p=0.8351 | F (9, 256) = 1.676, p=0.0949 |
| Panel C - Latency of repeated control rats | | | |
| Two-way mixed effects model | Session | Stage | Two-way interaction |
| | **F (2.746, 66.52) = 30.01, p<0.0001** | F (1, 25) = 2.092, p=0.1605 | F (9, 218) = 0.5312, p=0.8510 |
| Panel D - Latency of repeated stress rats | | | |
| Two-way mixed effects model | Session | Stage | Two-way interaction |
| | **F (2.654, 67.24) = 75.08, p<0.0001** | F (1, 27) = 0.1672, p=0.6859 | F (9, 228) = 0.6036, p=0.7934 |
| Panel E - Non-CS head entry rate of single control rats | | | |
| Two-way mixed effects model | Session | Stage | Two-way interaction |
| | F (1.799, 62.17) = 1.397, p=0.2544 | F (1, 35) = 0.1195, p=0.7317 | F (9, 311) = 0.7061, p=0.7033 |
| Panel F - Non-CS head entry rate of single stress rats | | | |
| Two-way mixed effects model | Session | Stage | Two-way interaction |
| | F (1.382, 39.30) = 3.049, p=0.0759 | F (1, 32) = 0.7595, p=0.3900 | F (9, 256) = 1.327, p=0.2230 |
| Panel G - Non-CS head entry rate of repeated control rats | | | |
| Two-way mixed effects model | Session | Stage | Two-way interaction |
| | F (1.551, 37.56) = 1.853, p=0.1775 | F (1, 25) = 0.09925, p=0.7553 | F (9, 218) = 0.7564, p=0.6568 |
| Panel H - Non-CS head entry rate of repeated stress rats | | | |
| Two-way mixed effects model | Session | Stage | Two-way interaction |
| | F (1.592, 40.33) = 2.486, p=0.1067 | F (1, 27) = 0.8976, p=0.3518 | F (9, 228) = 1.053, p=0.3986 |

| Supplementary Figure 7 | | | |
|---|---|---|---|
| Panel A - Experimental timeline | | | |
| Panel B - Extinction of non-estrus rats | | | |
| Three-way mixed-effects model | Session | Repeated treatment | Stress |
| | **F (1.701, 40.41) = 25.98, p<0.0001** | F (1, 26) = 1.006, p=0.3250 | F (1, 26) = 0.6605, p=0.4238 |
| | | Session X Repeated treatment | Session X Stress |
| | | **F (4, 95) = 2.474, p=0.0495** | F (4, 95) = 0.7064, p=0.5895 |
| | | Repeated treatment X Stress | Three-way interaction |
| | | F (1, 26) = 0.8120, p=0.3758 | F (4, 95) = 0.9246, p=0.4531 |
| Panel C - New learning of non-estrus rats | | | |
| Three-way mixed-effects model | Session | Repeated treatment | Stress |
| | **F (2.304, 54.72) = 11.38, p<0.0001** | **F (1, 26) = 12.02, p=0.0018** | **F (1, 26) = 8.958, p=0.0060** |
| | | Session X Repeated treatment | Session X Stress |
| | | F (4, 95) = 0.1924, p=0.9418 | F (4, 95) = 0.7302, p=0.5735 |
| | | Repeated treatment X Stress | Three-way interaction |
| | | F (1, 26) = 0.6856, p=0.4152 | F (4, 95) = 1.637, p=0.1713 |
| Panel D - Latency of non-estrus rats to food port after onset of extinction cue | | | |
| Three-way mixed-effects model | Session | Repeated treatment | Stress |
| | **F (2.681, 63.68) = 29.68, p<0.0001** | F (1, 26) = 3.741, p=0.0641 | F (1, 26) = 1.349, p=0.2560 |
| | | Session X Repeated treatment | Session X Stress |
| | | F (4, 95) = 0.7218, p=0.5792 | F (4, 95) = 1.285, p=0.2813 |
| | | Repeated treatment X Stress | Three-way interaction |
| | | F (1, 26) = 2.157, p=0.1539 | **F (4, 95) = 2.556, p=0.0438** |
| Panel E - Latency of non-estrus rats to food port after onset of new learning cue | | | |
| Three-way mixed-effects model | Session | Repeated treatment | Stress |
| | F (2.410, 57.24) = 2.147, p=0.1167 | **F (1, 26) = 4.359, p=0.0468** | F (1, 26) = 1.672, p=0.2074 |
| | | Session X Repeated treatment | Session X Stress |
| | | F (4, 95) = 1.928, p=0.1121 | F (4, 95) = 0.9314, p=0.4492 |
| | | Repeated treatment X Stress | Three-way interaction |
| | | F (1, 26) = 0.3145, p=0.5797 | F (4, 95) = 1.361, p=0.2534 |
| Panel F - Non-CS head entry rate of non-estrus rats | | | |
| Three-way mixed-effects model | Session | Repeated treatment | Stress |
| | **F (2.340, 55.58) = 13.93, p<0.0001** | F (1, 26) = 0.8347, p=0.3693 | F (1, 26) = 2.016, p=0.1675 |
| | | Session X Repeated treatment | Session X Stress |
| | | F (4, 95) = 1.126, p=0.3490 | F (4, 95) = 1.035, p=0.3933 |
| | | Repeated treatment X Stress | Three-way interaction |
| | | F (1, 26) = 1.633, p=0.2126 | F (4, 95) = 1.594, p=0.1822 |
| Panel G - Experimental timeline | | | |
| Panel H - Extinction of estrus rats | | | |
| Three-way mixed-effects model | Session | Repeated treatment | Stress |
| | **F (2.888, 67.14) = 27.64, p<0.0001** | F (1, 24) = 0.1182, p=0.7340 | F (1, 24) = 0.4462, p=0.5105 |
| | | Session X Repeated treatment | Session X Stress |
| | | **F (4, 93) = 3.955, p=0.0052** | F (4, 93) = 0.2375, p=0.9165 |
| | | Repeated treatment X Stress | Three-way interaction |
| | | F (1, 24) = 0.05671, p=0.8138 | F (4, 93) = 0.3745, p=0.8263 |
| Panel I - New learning of estrus rats | | | |
| Three-way mixed-effects model | Session | Repeated treatment | Stress |
| | **F (1.957, 45.49) = 9.901, p=0.0003** | **F (1, 24) = 8.092, p=0.0089** | F (1, 24) = 0.02198, p=0.8834 |
| | | Session X Repeated treatment | Session X Stress |
| | | F (4, 93) = 1.133, p=0.3457 | F (4, 93) = 0.7157, p=0.5833 |
| | | Repeated treatment X Stress | Three-way interaction |
| | | F (1, 24) = 3.643, p=0.0683 | F (4, 93) = 2.133, p=0.0829 |
| Panel J - Latency of estrus rats to food port after onset of extinction cue | | | |
| Three-way mixed-effects model | Session | Repeated treatment | Stress |
| | **F (2.397, 55.72) = 28.22, p<0.0001** | **F (1, 24) = 4.507, p=0.0443** | F (1, 24) = 0.06988 p=0.7938 |
| | | Session X Repeated treatment | Session X Stress |
| | | F (4, 93) = 1.593, p=0.1828 | F (4, 93) = 0.6261, p=0.6451 |
| | | Repeated treatment X Stress | Three-way interaction |
| | | F (1, 24) = 0.05082, p=0.8236 | **F (4, 93) = 2.620, p=0.0398** |
| Panel K - Latency of estrus rats to food port after onset of new learning cue | | | |
| Three-way mixed-effects model | Session | Repeated treatment | Stress |
| | **F (2.207, 51.31) = 8.775, p=0.0004** | **F (1, 24) = 7.812, p=0.0100** | F (1, 24) = 0.2579, p=0.6162 |
| | | Session X Repeated treatment | Session X Stress |
| | | F (4, 93) = 0.9955, p=0.4141 | F (4, 93) = 0.1994, p=0.9381 |
| | | Repeated treatment X Stress | Three-way interaction |
| | | F (1, 24) = 0.6698, p=0.4212 | F (4, 93) = 0.2633, p=0.9008 |
| Panel L - Non-CS head entry rate of estrus rats | | | |
| Three-way mixed-effects model | Session | Repeated treatment | Stress |
| | **F (1.860, 43.25) = 5.297, p=0.0101** | F (1, 24) = 3.479, p=0.0744 | F (1, 24) = 0.2258, p=0.6389 |
| | | Session X Repeated treatment | Session X Stress |
| | | **F (4, 93) = 4.602, p=0.0020** | F (4, 93) = 0.4347, p=0.7832 |
| | | Repeated treatment X Stress | Three-way interaction |
| | | F (1, 24) = 0.2734, p=0.6059 | F (4, 93) = 0.8502, p=0.4970 |

| Supplementary Figure 8 | | | |
|---|---|---|---|
| single stress non-estrus n=6; single stress estrus n=5; single control non-estrus n=8; single control estrus n=6; repeated stress non-estrus n=6; repeated stress estrus n=10; repeated control non-estrus n=8; repeated control estrus n=8 | | | |
| Panel A - Experimental timeline | | | |
| Panel B - Extinction of single control rats | | | |
| Two-way mixed effects model | Session | Stage | Two-way interaction |
| | **F (3.040, 35.73) = 16.60, p<0.0001** | F (1, 12) = 0.2328, p=0.6381 | F (4, 47) = 0.2366, p=0.9163 |
| Panel C - Extinction of single stress rats | | | |
| Two-way mixed effects model | Session | Stage | Two-way interaction |
| | **F (1.156, 9.826) = 5.339, p=0.0402** | F (1, 9) = 0.9106, p=0.3649 | F (4, 34) = 0.5771, p=0.6812 |
| Panel D - New learning of single control rats | | | |
| Two-way mixed effects model | Session | Stage | Two-way interaction |
| | **F (1.873, 22.01) = 4.938, p=0.0185** | F (1, 12) = 1.585, p=0.2320 | F (4, 47) = 0.4010, p=0.8069 |
| Panel E - New learning of single stress rats | | | |
| Two-way mixed effects model | Session | Stage | Two-way interaction |
| | **F (2.230, 18.96) = 8.522, p=0.0018** | **F (1, 9) = 5.905, p=0.0380** | F (4, 34) = 1.543, p=0.2118 |
| Post-hoc Tukey's multiple comparisons test | Session | 95.00% CI of diff. | Adjusted P Value |
| | 11 | -0.3944 to 0.1091 | 0.2264 |
| | 12 | -0.3237 to 0.05545 | 0.1434 |
| | 13 | -0.4844 to 0.1292 | 0.2131 |
| | **14** | **-0.5975 to -0.04892** | **0.0259** |
| | **15** | **-0.6712 to -0.04752** | **0.0289** |
| Panel F - Experimental timeline | | | |
| Panel G - Extinction of repeated control rats | | | |
| Two-way mixed effects model | Session | Stage | Two-way interaction |
| | **F (2.257, 30.47) = 23.16, p<0.0001** | F (1, 15) = 1.099, p=0.3110 | F (4, 54) = 1.426, p=0.2380 |
| Panel H - Extinction of repeated stress rats | | | |
| Two-way mixed effects model | Session | Stage | Two-way interaction |
| | **F (2.150, 28.49) = 16.62, p<0.0001** | F (1, 14) = 1.673, p=0.2168 | F (4, 53) = 0.7550, p=0.5592 |
| Panel I - New learning of repeated control rats | | | |
| Two-way mixed effects model | Session | Stage | Two-way interaction |
| | **F (2.098, 28.32) = 3.997, p=0.0279** | F (1, 15) = 0.1163, p=0.7379 | F (4, 54) = 1.038, p=0.3961 |
| Panel J - New learning of repeated stress rats | | | |
| Two-way mixed effects model | Session | Stage | Two-way interaction |
| | **F (2.179, 28.87) = 8.393, p=0.0010** | F (1, 14) = 0.04246, p=0.8397 | F (4, 53) = 1.614, p=0.1845 |

| Supplementary Figure 9 | | | |
|---|---|---|---|
| single control non-estrus n=8; single control estrus n=6; repeated control non-estrus n=8; repeated control estrus n=8 | | | |
| Panel A - Experimental timeline | | | |
| Panel B - Extinction of control non-estrus rats | | | |
| Two-way mixed effects model | Session | Repeated treatment | Two-way interaction |
| | **F (1.574, 22.83) = 18.66, p<0.0001** | F (1, 16) = 3.488, p=0.0802 | **F (4, 58) = 3.349, p=0.0156** |
| Post-hoc Tukey's multiple comparisons test | Session | 95.00% CI of diff. | Adjusted P Value |
| | 11 | Not applicable, as S11 is normalized to 1 | |
| | 12 | -0.2095 to 0.1789 | 0.8681 |
| | 13 | -0.06278 to 0.6952 | 0.0956 |
| | 14 | -0.07438 to 0.6304 | 0.1119 |
| | 15 | -0.1868 to 1.024 | 0.1371 |
| Panel C- Extinction of control estrus rats | | | |
| Two-way mixed effects model | Session | Repeated treatment | Two-way interaction |
| | **F (2.127, 22.86) = 22.97, p<0.0001** | F (1, 11) = 0.3153, p=0.5857 | F (4, 43) = 1.464, p=0.2297 |
| Panel D - New learning of control non-estrus rats | | | |
| Two-way mixed effects model | Session | Repeated treatment | Two-way interaction |
| | **F (2.777, 40.26) = 6.490, p=0.0014** | **F (1, 16) = 5.616, p=0.0307** | F (4, 58) = 2.008, p=0.1053 |
| Post-hoc Tukey's multiple comparisons test | Session | 95.00% CI of diff. | Adjusted P Value |
| | **11** | **-0.3664 to -0.007950** | **0.0417** |
| | 12 | -0.3770 to 0.01978 | 0.0744 |
| | 13 | -0.2780 to 0.1612 | 0.5783 |
| | 14 | -0.5227 to 0.07314 | 0.1293 |
| | **15** | **-0.8463 to -0.05611** | **0.0305** |
| Panel E - New learning of control estrus rats | | | |
| Two-way mixed effects model | Session | Repeated treatment | Two-way interaction |
| | F (1.519, 16.33) = 2.330, p=0.1377 | F (1, 11) = 0.4138, p=0.5332 | F (4, 43) = 0.6175, p=0.6524 |

| Supplementary Figure 10 | | | |
|---|---|---|---|
| single stress non-estrus n=6; single stress estrus n=5; single control non-estrus n=8; single control estrus n=6; repeated stress non-estrus n=6; repeated stress estrus n=10; repeated control non-estrus n=8; repeated control estrus n=8 | | | |
| Panel A - Latency of single treatment non-estrus rats to food port after onset of white noise (extinction) cue | | | |
| Two-way mixed effects model | Session | Stress | Two-way interaction |
| | **F (2.450, 28.18) = 10.62, p=0.0002** | F (1, 12) = 0.03231, p=0.8603 | F (4, 46) = 0.5770, p=0.6807 |
| Panel B - Latency of single treatment estrus rats to food port after onset of white noise (extinction) cue | | | |
| Two-way mixed effects model | Session | Stress | Two-way interaction |
| | **F (2.152, 18.83) = 9.262, p=0.0013** | F (1, 9) = 0.07972, p=0.7841 | F (4, 35) = 1.581, p=0.2009 |
| Panel C - Latency of single treatment non-estrus rats to food port after onset of tone (new learning) cue | | | |
| Two-way mixed effects model | Session | Stress | Two-way interaction |
| | F (1.777, 20.44) = 2.973, p=0.0787 | F (1, 12) = 0.1840, p=0.6756 | F (4, 46) = 0.9952, p=0.4198 |
| Panel D - Latency of single treatment estrus rats to food port after onset of tone (new learning) cue | | | |
| Two-way mixed effects model | Session | Stress | Two-way interaction |
| | **F (1.457, 12.75) = 4.626, p=0.0400** | F (1, 9) = 0.03249, p=0.8609 | F (4, 35) = 0.03032, p=0.9981 |
| Panel E - Non-CS head entry rate of single treatment non-estrus rats | | | |
| Two-way mixed effects model | Session | Stress | Two-way interaction |
| | **F (1.288, 14.81) = 4.948, p=0.0346** | F (1, 12) = 0.007087, p=0.9343 | F (4, 46) = 0.4561, p=0.7674 |
| Panel F - Non-CS head entry rate of single treatment estrus rats | | | |
| Two-way mixed effects model | Session | Stress | Two-way interaction |
| | F (1.873, 16.39) = 2.864, p=0.0885 | F (1, 9) = 0.004967, p=0.9454 | F (4, 35) = 0.7149, p=0.5874 |
| Panel G - Latency of repeated treatment non-estrus rats to food port after onset of white noise (extinction) cue | | | |
| Two-way mixed effects model | Session | Stress | Two-way interaction |
| | **F (2.718, 33.30) = 20.72, p<0.0001** | **F (1, 14) = 5.832, p=0.0300** | **F (4, 49) = 3.306, p=0.0178** |
| Post-hoc Tukey's multiple comparisons test | Session | 95.00% CI of diff. | Adjusted P Value |
| | 11 | -0.3875 to 0.7125 | 0.5155 |
| | 12 | -0.5067 to 0.6798 | 0.7571 |
| | 13 | -0.05530 to 1.528 | 0.0657 |
| | 14 | -0.3118 to 1.893 | 0.1408 |
| | **15** | **0.2795 to 2.811** | **0.0283** |
| Panel H - Latency of repeated treatment estrus rats to food port after onset of white noise (extinction) cue | | | |
| Two-way mixed effects model | Session | Stress | Two-way interaction |
| | **F (2.209, 32.03) = 23.42, p<0.0001** | F (1, 15) = 0.001132, p=0.9736 | F (4, 58) = 1.760, p=0.1493 |
| Panel I - Latency of repeated treatment non-estrus rats to food port after onset of tone (new learning) cue | | | |
| Two-way mixed effects model | Session | Stress | Two-way interaction |
| | F (2.532, 31.01) = 0.5048, p=0.6513 | F (1, 14) = 2.734, p=0.1205 | F (4, 49) = 1.234, p=0.3089 |
| Panel J - Latency of repeated treatment estrus rats to food port after onset of tone (new learning) cue | | | |
| Two-way mixed effects model | Session | Stress | Two-way interaction |
| | **F (2.495, 36.17) = 4.484, p=0.0126** | F (1, 15) = 1.284, p=0.2749 | F (4, 58) = 0.5918, p=0.6699 |
| Panel K - Non-CS head entry rate of repeated treatment non-estrus rats | | | |
| Two-way mixed effects model | Session | Stress | Two-way interaction |
| | **F (2.132, 26.12) = 9.351, p=0.0007** | **F (1, 14) = 5.481, p=0.0345** | F (4, 49) = 1.804, p=0.1431 |
| Post-hoc Tukey's multiple comparisons test | Session | 95.00% CI of diff. | Adjusted P Value |
| | 11 | -584.0 to 6.814 | 0.0541 |
| | 12 | -388.4 to 21.44 | 0.0738 |
| | 13 | -375.1 to 25.86 | 0.0808 |
| | 14 | -389.1 to 63.52 | 0.1296 |
| | 15 | -409.6 to 35.70 | 0.0825 |
| Panel L - Non-CS head entry rate of repeated treatment estrus rats | | | |
| Two-way mixed effects model | Session | Stress | Two-way interaction |
| | **F (1.591, 23.07) = 7.820, p=0.0043** | F (1, 15) = 0.4197, p=0.5269 | F (4, 58) = 0.8965, 0.4720 |
| Panel M - Latency of stressed non-estrus rats to food port after onset of white noise (extinction) cue | | | |
| Two-way mixed effects model | Session | Repeated treatment | Two-way interaction |
| | **F (2.274, 21.03) = 8.928, p=0.0011** | **F (1, 10) = 5.660, p=0.0387** | F (4, 37) = 0.3862, p=0.8171 |
| Post-hoc Tukey's multiple comparisons test | Session | 95.00% CI of diff. | Adjusted P Value |
| | 11 | -0.1200 to 1.280 | 0.0945 |
| | **12** | **0.03618 to 1.340** | **0.0405** |
| | 13 | -0.2333 to 2.183 | 0.0973 |
| | 14 | -0.4927 to 2.262 | 0.1824 |
| | 15 | -0.5295 to 2.112 | 0.1995 |
| Panel N - Latency of stressed estrus rats to food port after onset of white noise (extinction) cue | | | |
| Two-way mixed effects model | Session | Repeated treatment | Two-way interaction |
| | **F (2.039, 25.49) = 13.05, p=0.0001** | F (1, 13) = 3.704, p=0.0765 | **F (4, 50) = 2.669, p=0.0428** |
| Post-hoc Tukey's multiple comparisons test | Session | 95.00% CI of diff. | Adjusted P Value |
| | 11 | -0.4499 to 2.615 | 0.1243 |
| | 12 | -0.1114 to 2.607 | 0.0652 |
| | 13 | -0.1816 to 2.420 | 0.08 |
| | 14 | -1.015 to 1.851 | 0.5092 |
| | 15 | -2.106 to 2.206 | 0.9511 |
| Panel O - Latency of stressed non-estrus rats to food port after onset of tone (new learning) cue | | | |
| Two-way mixed effects model | Session | Repeated treatment | Two-way interaction |
| | F (1.401, 12.96) = 2.927, p=0.1018 | **F (1, 10) = 9.633, p=0.0112** | **F (4, 37) = 2.917, p=0.0341** |
| Post-hoc Tukey's multiple comparisons test | Session | 95.00% CI of diff. | Adjusted P Value |
| | **11** | **0.4897 to 2.128** | **0.0052** |
| | **12** | **0.4163 to 1.944** | **0.0086** |
| | 13 | -0.4443 to 1.773 | 0.1802 |
| | **14** | **0.07642 to 1.607** | **0.0355** |
| | 15 | -0.6200 to 1.167 | 0.487 |
| Panel P - Latency of stressed estrus rats to food port after onset of tone (new learning) cue | | | |
| Two-way mixed effects model | Session | Repeated treatment | Two-way interaction |
| | **F (1.918, 23.97) = 8.501, p=0.0018** | **F (1, 13) = 9.236, p=0.0095** | F (4, 50) = 0.7922, p=0.5358 |
| Post-hoc Tukey's multiple comparisons test | Session | 95.00% CI of diff. | Adjusted P Value |
| | 11 | -0.1238 to 2.598 | 0.0694 |
| | **12** | **0.1635 to 3.081** | **0.0352** |
| | **13** | **0.005732 to 2.663** | **0.0493** |
| | 14 | -0.2082 to 2.555 | 0.0793 |
| | 15 | -0.8024 to 2.987 | 0.171 |
| Panel Q - Non-CS head entry rate of stressed non-estrus rats | | | |
| Two-way mixed effects model | Session | Stress | Two-way interaction |
| | **F (1.846, 17.07) = 5.970, p=0.0122** | F (1, 10) = 1.599, p=0.2348 | F (4, 37) = 1.530, p=0.2137 |
| Panel R - Non-CS head entry rate of stressed estrus rats | | | |
| Two-way mixed effects model | Session | Stress | Two-way interaction |
| | **F (1.598, 19.98) = 5.409, p=0.0181** | F (1, 13) = 4.392, p=0.0562 | **F (4, 50) = 4.701, p=0.0027** |
| Post-hoc Tukey's multiple comparisons test | Session | 95.00% CI of diff. | Adjusted P Value |
| | 11 | -390.1 to 13.50 | 0.0632 |
| | **12** | **-443.2 to -14.84** | **0.0387** |
| | **13** | **-266.2 to -48.40** | **0.0091** |
| | 14 | -217.3 to 223.7 | 0.9713 |
| | 15 | -268.7 to 136.0 | 0.4061 |

| Supplementary Figure 11 | | | |
|---|---|---|---|
| single stress non-estrus n=10; single stress estrus n=3; single control non-estrus n=11; single control estrus n=12; repeated stress non-estrus n=13; repeated stress estrus n=7; repeated control non-estrus n=20; repeated control estrus n=7 | | | |
| Panel A - Percent time in open arms | | | |
| One way ANOVA | Treatment | | |
| | F (7, 75) = 0.5688, p=0.7790 | | |
| Tukey's multiple comparisons test | | | |
| Comparison | Mean Diff. | 95.00% CI of diff. | Adjusted P Value |
| Single control estrus V Single control non-estrus | 8.338 | -12.93 to 29.61 | 0.9227 |
| Single control estrus V Single stress estrus | -0.9329 | -33.82 to 31.96 | >0.9999 |
| Single control estrus V Single stress non-estrus | 4.458 | -17.36 to 26.27 | 0.9982 |
| Single control estrus V Repeated control estrus | 8.145 | -16.09 to 32.38 | 0.9652 |
| Single control estrus V Repeated control non-estrus | 4.473 | -14.13 to 23.08 | 0.995 |
| Single control estrus V Repeated stress estrus | -0.2884 | -24.52 to 23.94 | >0.9999 |
| Single control estrus V Repeated stress non-estrus | -1.563 | -21.96 to 18.83 | >0.9999 |
| Single control non-estrus V Single stress estrus | -9.271 | -42.46 to 23.92 | 0.9878 |
| Single control non-estrus V Single stress non-estrus | -3.879 | -26.14 to 18.38 | 0.9994 |
| Single control non-estrus V Repeated control estrus | -0.1927 | -24.83 to 24.44 | >0.9999 |
| Single control non-estrus V Repeated control non-estrus | -3.864 | -22.99 to 15.26 | 0.9983 |
| Single control non-estrus V Repeated stress estrus | -8.626 | -33.26 to 16.01 | 0.9566 |
| Single control non-estrus V Repeated stress non-estrus | -9.9 | -30.77 to 10.97 | 0.8161 |
| Single stress estrus V Single stress non-estrus | 5.391 | -28.15 to 38.93 | 0.9996 |
| Single stress estrus V Repeated control estrus | 9.078 | -26.08 to 44.24 | 0.9923 |
| Single stress estrus V Repeated control non-estrus | 5.406 | -26.14 to 36.95 | 0.9994 |
| Single stress estrus V Repeated stress estrus | 0.6444 | -34.52 to 35.81 | >0.9999 |
| Single stress estrus V Repeated stress non-estrus | -0.6299 | -33.27 to 32.01 | >0.9999 |
| Single stress non-estrus V Repeated control estrus | 3.687 | -21.42 to 28.80 | 0.9998 |
| Single stress non-estrus V Repeated control non-estrus | 0.015 | -19.72 to 19.75 | >0.9999 |
| Single stress non-estrus V Repeated stress estrus | -4.747 | -29.86 to 20.36 | 0.9989 |
| Single stress non-estrus V Repeated stress non-estrus | -6.021 | -27.45 to 15.41 | 0.9874 |
| Repeated control estrus V Repeated control non-estrus | -3.672 | -26.05 to 18.70 | 0.9996 |
| Repeated control estrus V Repeated stress estrus | -8.433 | -35.67 to 18.80 | 0.9779 |
| Repeated control estrus V Repeated stress non-estrus | -9.708 | -33.59 to 14.18 | 0.9079 |
| Repeated control non-estrus V Repeated stress estrus | -4.762 | -27.14 to 17.61 | 0.9977 |
| Repeated control non-estrus V Repeated stress non-estrus | -6.036 | -24.19 to 12.12 | 0.9671 |
| Repeated stress estrus V Repeated stress non-estrus | -1.274 | -25.16 to 22.61 | >0.9999 |
| Panel B - Total distance travelled | | | |
| One way ANOVA | Treatment | | |
| | F (7, 75) = 0.1925, p=0.9862 | | |
| Tukey's multiple comparisons test | | | |
| Comparison | Mean Diff. | 95.00% CI of diff. | Adjusted P Value |
| Single control estrus V Single control non-estrus | 0.5351 | -10.00 to 11.07 | >0.9999 |
| Single control estrus V Single stress estrus | -1.075 | -17.37 to 15.22 | >0.9999 |
| Single control estrus V Single stress non-estrus | 1.74 | -9.068 to 12.55 | 0.9996 |
| Single control estrus V Repeated control estrus | 0.1279 | -11.88 to 12.13 | >0.9999 |
| Single control estrus V Repeated control non-estrus | -1.255 | -10.47 to 7.962 | 0.9999 |
| Single control estrus V Repeated stress estrus | -0.03027 | -12.03 to 11.97 | >0.9999 |
| Single control estrus V Repeated stress non-estrus | -1.416 | -11.52 to 8.689 | 0.9998 |
| Single control non-estrus V Single stress estrus | -1.61 | -18.05 to 14.83 | >0.9999 |
| Single control non-estrus V Single stress non-estrus | 1.205 | -9.824 to 12.23 | >0.9999 |
| Single control non-estrus V Repeated control estrus | -0.4073 | -12.61 to 11.80 | >0.9999 |
| Single control non-estrus V Repeated control non-estrus | -1.79 | -11.26 to 7.685 | 0.9989 |
| Single control non-estrus V Repeated stress estrus | -0.5654 | -12.77 to 11.64 | >0.9999 |
| Single control non-estrus V Repeated stress non-estrus | -1.951 | -12.29 to 8.390 | 0.9989 |
| Single stress estrus V Single stress non-estrus | 2.815 | -13.80 to 19.43 | 0.9995 |
| Single stress estrus V Repeated control estrus | 1.203 | -16.22 to 18.62 | >0.9999 |
| Single stress estrus V Repeated control non-estrus | -0.1795 | -15.81 to 15.45 | >0.9999 |
| Single stress estrus V Repeated stress estrus | 1.045 | -16.37 to 18.46 | >0.9999 |
| Single stress estrus V Repeated stress non-estrus | -0.3406 | -16.51 to 15.83 | >0.9999 |
| Single stress non-estrus V Repeated control estrus | -1.612 | -14.05 to 10.83 | >0.9999 |
| Single stress non-estrus V Repeated control non-estrus | -2.995 | -12.77 to 6.781 | 0.9792 |
| Single stress non-estrus V Repeated stress estrus | -1.77 | -14.21 to 10.67 | 0.9998 |
| Single stress non-estrus V Repeated stress non-estrus | -3.156 | -13.77 to 7.461 | 0.9825 |
| Repeated control estrus V Repeated control non-estrus | -1.382 | -12.47 to 9.702 | >0.9999 |
| Repeated control estrus V Repeated stress estrus | -0.1581 | -13.65 to 13.33 | >0.9999 |
| Repeated control estrus V Repeated stress non-estrus | -1.544 | -13.38 to 10.29 | >0.9999 |
| Repeated control non-estrus V Repeated stress estrus | 1.224 | -9.861 to 12.31 | >0.9999 |
| Repeated control non-estrus V Repeated stress non-estrus | -0.1612 | -9.154 to 8.831 | >0.9999 |
| Repeated stress estrus V Repeated stress non-estrus | -1.385 | -13.22 to 10.45 | >0.9999 |
